# Single-cell methylation sequencing data reveal succinct metastatic migration histories and tumor progression models

**DOI:** 10.1101/2021.03.22.436475

**Authors:** Yuelin Liu, Xuan Cindy Li, Farid Rashidi Mehrabadi, Alejandro A. Schäffer, Drew Pratt, David R. Crawford, Salem Malikić, Erin K. Molloy, Vishaka Gopalan, Stephen M. Mount, Eytan Ruppin, Kenneth Aldape, S. Cenk Sahinalp

**Author notes:** these authors contributed equally to this work (Yuelin Liu, Xuan Cindy Li).

## Abstract

Recent studies exploring the impact of methylation in tumor evolution suggest that while the methylation status of many of the CpG sites are preserved across distinct lineages, others are altered as the cancer progresses. Since changes in methylation status of a CpG site may be retained in mitosis, they could be used to infer the progression history of a tumor via single-cell lineage tree reconstruction. In this work, we introduce the first principled distance-based computational method, `Sgootr`, for inferring a tumor’s single-cell methylation lineage tree and jointly identifying lineage-informative CpG sites which harbor changes in methylation status that are retained along the lineage. We apply `Sgootr` on the single-cell bisulfite-treated whole genome sequencing data of multiregionally-sampled tumor cells from 9 metastatic colorectal cancer patients made available by Bian *et al*., as well as multiregionally-sampled single-cell reduced-representation bisulfite sequencing data from a glioblastoma patient made available by Chaligne *et al*.. We demonstrate that the tumor lineages constructed reveal a simple model underlying colorectal tumor progression and metastatic seeding. A comparison of `Sgootr` against alternative approaches shows that `Sgootr` can construct lineage trees with fewer migration events and more in concordance with the sequential-progression model of tumor evolution, in time a fraction of that used in prior studies. Interestingly, lineage-informative CpG sites identified by `Sgootr` are in inter-CpG island (CGI) regions, as opposed to CGI’s, which have been the main regions of interest in genomic methylation-related analyses. `Sgootr` is implemented as a `Snakemake` workflow, available at https://github.com/algo-cancer/Sgootr.

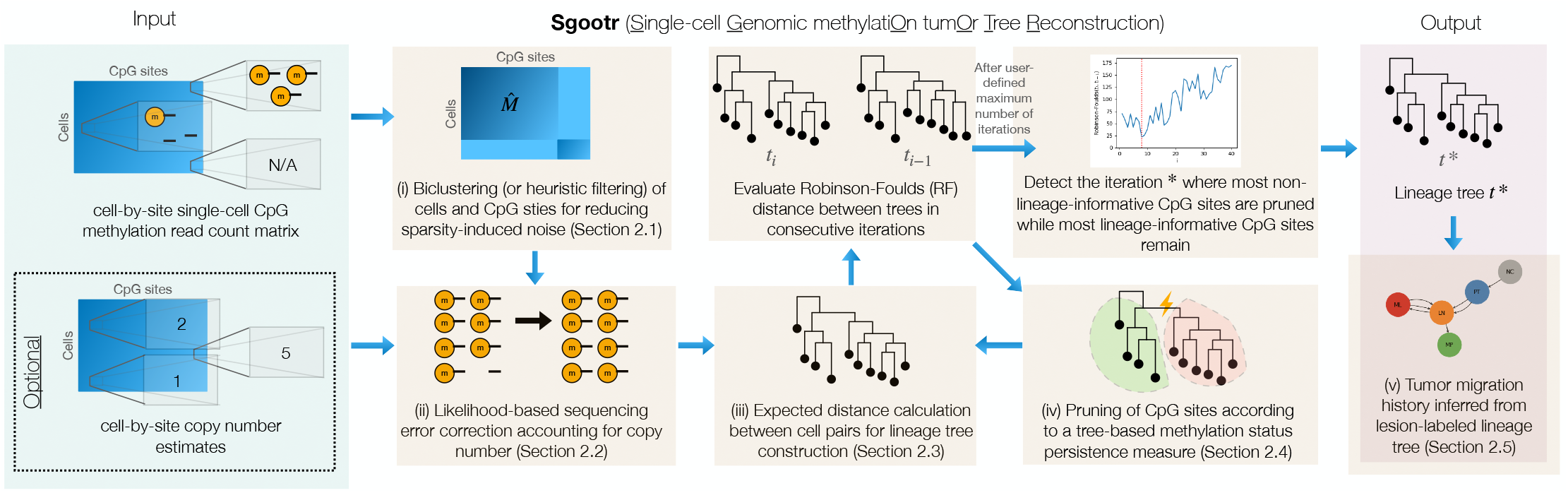

## 1 Introduction

The recent rise of single-cell sequencing technology empowers more accurate tumor lineage inference by allowing the examination of intratumor heterogeneity at a cellular resolution. However, since single-cell sequencing data is derived from an incredibly limited amount of genetic material, the signals obtained are more scarce and unstable than those from bulk sequencing [23]. DNA methylation, the addition of a methyl group to cytosine, which results in the formation of 5-methylcytosine (5mC) especially in the context of CpG sites, is an epigenetic marker that has been extensively studied for its role in regulating gene expression and maintaining cellular memory [20]. Prior research suggests the changes in methylation status at CpG sites in cancer cells may provide a greater amount of observable evidence for tumor evolution than single-nucleotide variations (SNVs) or copy number aberrations (CNAs) [38, 3].

Recent studies have leveraged CpG methylation in single cells sampled from colorectal cancer (CRC) [2], chronic lymphocitic leukemia (CLL) [14], glioblastoma (GBM) and IDH-mutant glioma [5] tumors to construct lineages, showing CpG methylation to be a valuable signal for lineage inference. However, further examination reveals that data obtained via single-cell bisulfite sequencing (scBS-seq) [2], single-cell reduced representation bisulfite sequencing (scRRBS)[15, 16], or multiplexed scRRBS (MscRRBS) [14] protocols exhibit high level of sparsity. Not only that cells with subpar bisulfite conversion rates have information at few CpG sites, of the millions of unique CpG sites across all cells in a dataset, less than one out of a hundred are sequenced in a sufficient number of cells to be useful in lineage reconstruction; furthermore, even when a CpG site is sequenced in a cell, it is oftentimes covered by less than a few sequencing reads, which is not likely to capture heterozygosity in aneuploid cancers.

Besides sparsity, another key challenge is that not all CpG sites have their methylation statuses stably retained during in tumor evolution. In particular, Meir *et al*. posited two models of methylation dynamics: *mixture*, where the methylation status of a CpG site is resampled (from the parental status) in each cell replication, and *persistency*, where that is an exact copy of the parent cell [28]. It is the CpG sites whose status follows the persistency model would be informative in tumor lineage reconstruction because the maintenance of information from the parental generation is the necessary condition for the infinite sites assumption [21, 27] and Dollo’s parsimony [8], which form the basis for tumor lineage inference tools based on mutation profiles. However, as of now, there is no known set of CpG sites that follow either inheritance model, highlighting the necessity to perform intentional CpG site selection while reconstructing tumor lineages by CpG methylation. In this work, we introduce `Sgootr` (Single-cell Genomic methylatiOn tumOr Tree Reconstruction), the first distance-based computational method to jointly select informative CpG sites and reconstruct tumor lineages from single-cell methylation data. Application of `Sgootr` to a multiregionally-sampled metastatic CRC scBS-seq dataset[2] and a multiregionally-sampled GBM MscRRBS patient sample [5] reveals tumor progression models and metastatic migration histories simpler than previously reported. A comparison of `Sgootr` against alternative lineage-reconstruction and site-selection approaches on the same datasets shows that it infers a simpler migration history in a shorter amount of time. Interestingly lineage-informative CpG sites identified by `Sgootr` appear to be primarily in inter-CpG island (CGI) regions, as opposed to CGI’s which have been the main regions of interest in genomic methylation-related analyses.

## 2 Methods

`Sgootr` consists of five key components: (i) biclustering of cells and sites for reducing sparsity-induced noise (Section 2.1); (ii) likelihood-based sequencing error correction accounting for copy number variation (Section 2.2); (iii) expected distance calculation between cell pairs for tree construction (Section 2.3); (iv) pruning of CpG sites according to a tree-based methylation status persistence measure (Section 2.4); and (v) inference of migration history from the lesion-labeled tree (Section 2.5). Note that components (iii) and (iv) are iteratively applied. Code for this work is available on GitHub as a `Snakemake` [29] workflow. Results from this work are reproducible, and intermediate experiment outputs of `Sgootr` are available for further analysis.

### 2.1 Biclustering of cells and sites for reducing sparsity-induced noise

`Sgootr` requires a fairly accurate estimate of the differences between the methylation status of distinct CpG sites across all cell pairs. This necessitates filtering out those cells for which the number of CpG sites with non-zero read coverage is low and CpG sites with non-zero coverage in only a few cells. Ideally, we would like to coordinate the reductions in cells and CpG sites so as to minimize information loss, which can be achieved via an integer linear program (ILP)-based biclustering formulation as follows.

Let the site coverage data be represented in a cell-by-site matrix *M*_*v*×*u*_, where *v* is the number of cells and *u* is the number of CpG sites; *m*_ij_ = 1 if site *j* is covered (by at least one read) in cell *i*, and *m*_ij_ = − 1 otherwise. Let *α* and *β* respectively be the specified fraction of cells and sites to be kept. Given *M, α*, and *β*, we wish to compute a biclustering - namely, a selection of rows of columns - of *M*, 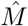, so that we maximize the number of CpG sites covered in the resulting ⌊*αv*⌋×⌊*βu*⌋ submatrix in which rows and columns respectively correspond to the selected cells and sites.

To solve this biclustering problem, let *C* ∈ {0, 1} ^*v*^, *S* ∈ {0, 1} ^*u*^ be (unknown) binary vectors respectively indicating whether a cell or site is kept. In addition, let *A* ∈ {0, 1} ^*v*×*u*^ denote (unknown) binary matrix such that *a*_*ij*_ = 1 if and only if *c*_*i*_ = 1 and *s*_*j*_ = 1 (i.e., if cell *i* and site *j* are both kept). The ILP formulation is:

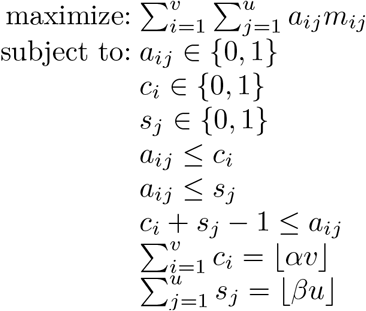

The output submatrix 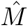 is obtained by taking cells {*i*|*c*_*i*_ = 1} and sites {*j*|*s*_*j*_ = 1} from *M*.

Our implementation of the above formulation employs the Gurobi ILP solver, a licensed software which is free to use for academic purposes [17]. The performance of our implementation with respect run time, memory and the accuracy (i.e. optimality gap) with respect to the input size and other parameters, as well as a strategy on how to set *α* and *β* are explored in detail in Supplementary Section S1. For problem instances in which our implementation does not lead to a near-optimal solution within reasonable time/memory (with respect to the optimality gap), or if the Gurobi ILP solver is unavailable, we apply the following heuristic filters to the cells and sites in the dataset at hand. (i) Among all input cells, remove low-quality ones with coverage in fewer than 4 million CpG sites; (ii) remove CpG sites covered in less than 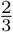 of the remaining cells. In addition to the (ILP based or heuristic) biclustering, it is advisable to remove CpG sites on sex chromosomes, which may confound the findings, and chromosome 21 peri-centromeric regions (chr21:9825000-9828000), which we discovered to be an overlooked alignment artifact.

After filtering the input for reducing sparsity, we perform sequencing error correction as described in Section 2.2. Since this may further introduce sparsity, in an additional post error correction step, we may remove sites covering 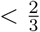 of cells, and then remove cells with coverage in 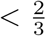 of the remaining CpG sites.

Overall, our filtering approach aims to make sure that the cells remaining are well-covered, and that each cell pair have at least 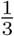 of the selected CpG sites as shared dimensions along which cell-to-cell distance can be measured.

### 2.2 Likelihood-based sequencing error correction accounting for copy number

Errors in sequencing may provide false evidence for an underlying allele methylation status different from the truth. We present a maximum log-likelihood approach to correct likely errors in methylation reads, incorporating site copy number estimates in each cell (Algorithm 1). For a CpG site in a cell, given *n* ≥ 0 reads with a methylated status, *m* ≥ 0 reads with an unmethylated status, its copy number *c*, and sequencing error probability 0 ≤ *ϵ* ≤ 1 (which we set to .01 in our experiments to proxy the per-base sequencing error rate in Illumina sequencing instruments), we can enumerate the likelihood of drawing the observed reads from all possible underlying allele statuses, letting 0 ≤ *γ* ≤ *c* be the number of alleles with the CpG site methylated. Note that in Algorithm 1, 1^c^ and 0^c^ respectively denote the events that all *c* alleles are methylated and unmethylated; 1^γ^0^c−γ^ denotes the event where *γ* alleles are methylated and *c* − *γ* are not.

Sequencing error correction takes place when we have *n >* 0 and *m >* 0, but the allele status with the highest log-likelihood is homozygous. In that case, we correct the reads with the minority methylation status to the majority one. When *c* = 1 the site will implicitly be identified as homozygous and the methylation status of any read that does not conform to the majority allele will be corrected. In the case where sequencing error is needed, yet *n* = *m*, which will likely happen for *c* = 1, the CpG site is discarded. In case copy number information is not available, one can assume an appropriate (uninformative) copy number for all sites in all cells (e.g. *c* = 2 for diploid). Sex chromosomes are always excluded from our analysis. We use the sequence error-corrected reads for pairwise distance estimation between cells, as described in the following section.

#### Algorithm 1 Sequencing error correction accounting for copy number

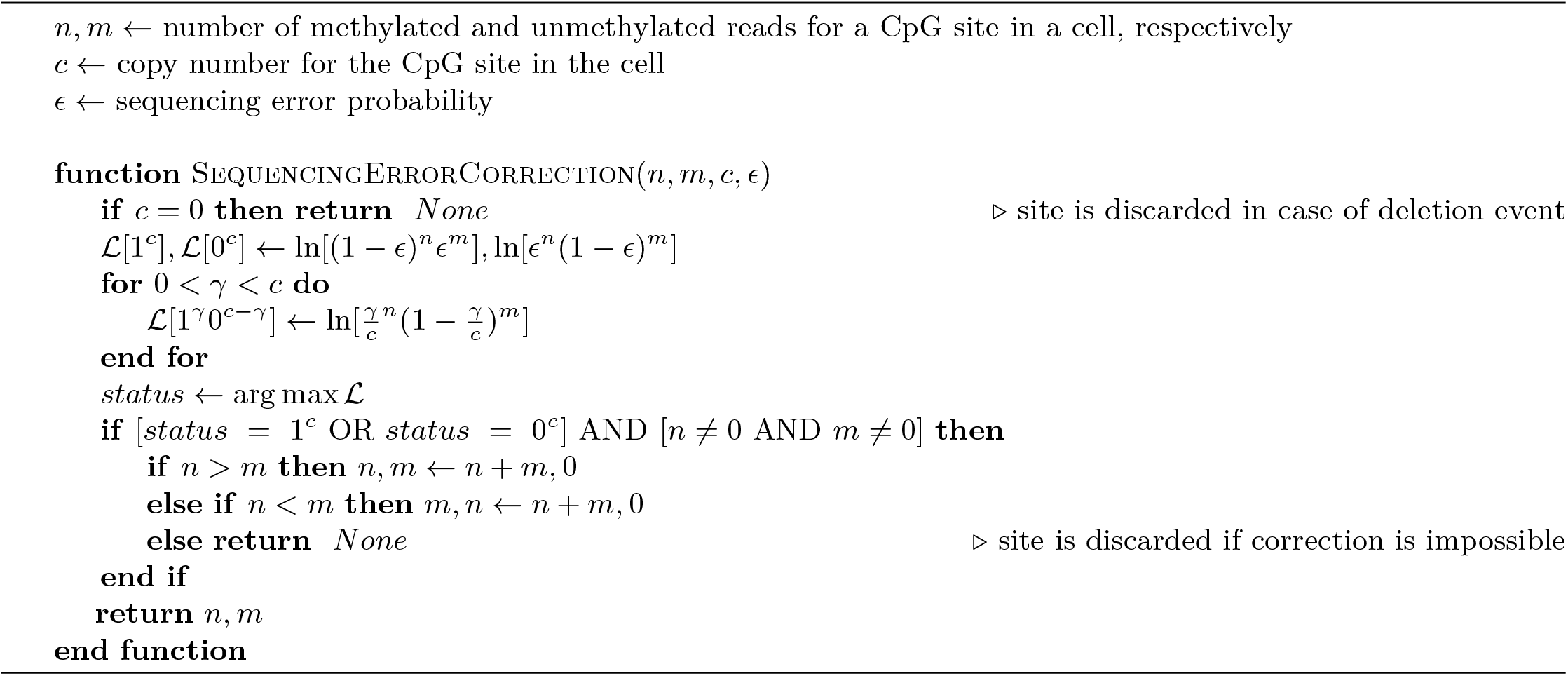

### 2.3 Expected distance calculation between cell pairs for tree construction

In this section, we present a formulation for computing the expected distance between two cells given their respective copy number and (sequencing error-corrected) reads. We consider such a formulation with the goal of correcting for the potential bias contributed by low coverage CpG sites. Intuitively, this distance measure serves as an estimate for the expected proportion of methylation status altered alleles across all CpG sites between a cell pair.

Given a cell, let the copy number at a CpG site be *c*, and we have 0 ≤ *γ* ≤ *c* alleles with the CpG site methylated, and *c* − *γ* alleles with the CpG site unmethylated. To allow modeling preferential allele sampling based on site methylation status as previously observed [31], we additionally introduce parameter *p*, probability of drawing a read from the allele with the CpG site methylated given a pair of alleles heterozygous for the site. We set *p* to the uninformative .5 in our experiments (see Supplementary Section S6 for when to choose alternative values). Then, we can compute the probability of drawing from an allele with the CpG site methylated by normalizing over all alleles (Equation 1):

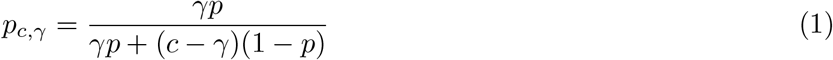

It follows that the probability of drawing from an allele with the CpG site unmethylated is 1 − *p*_c,γ_.

Consider the case where we observe *n* reads at a CpG site for a cell, and all *n* reads are methylated for the site. Let the copy number at the site for the cell be *c*, the probability of sampling all *n* methylated reads from *c* alleles with the CpG site methylated (i.e. *γ* = *c*) is: P(reads | 1^c^) = 1; here 1^c^ is the event that in all *c* copies the CpG site is methylated, and reads is the event that all *n* sampled reads are methylated. The probability of drawing all *n* methylated reads from *c* alleles with the CpG site unmethylated (i.e. *γ* = 0) is: P(reads | 0^c^) = 0. To compute the probability of drawing *n* methylated reads from *c* alleles with mixed methylation status for the site, we need to sum the probabilities over all other possible values of *γ*, the number of alleles with the site methylated, assuming that all other possible values of *γ* are equally likely; and when there is a copy number loss (i.e. *c* = 1), the formulation assigns only non-zero probability to a homozygous combined allele status:

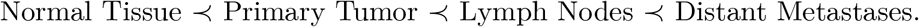

Given user-defined prior probabilities P(1^c^), P(0^c^), and P(mixed), which we all set to the uninformative .33 in our experiments (see Supplementary Section S6 for when to choose alternative values), let *a* = P(reads | 1^c^)P(1^c^) + P(reads | mixed)P(mixed) + P(reads | 0^c^)P(0^c^), then we apply Bayes’ Theorem to get the likelihood of any allele status given the observed reads (Equations 2):

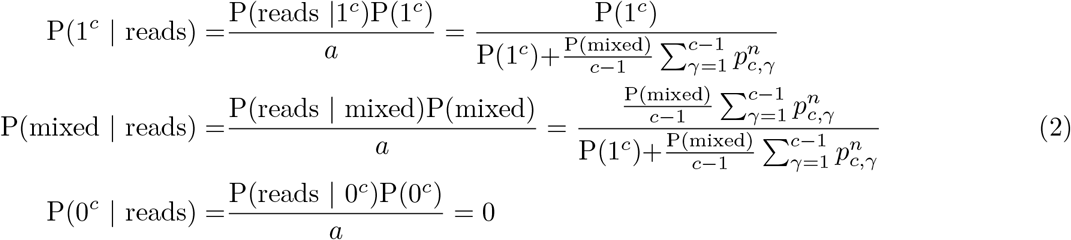

In the case where the *n* reads all have the CpG site unmethylated, the three values can be computed similarly with P(reads | 1^c^) = 0 and P(reads | 0^c^) = 1. In the case we observe among the *n* reads both ones with the CpG site methylated and ones with the site unmethylated, we have P(mixed reads)=1, and P(1^c^ reads) = P(0^c^ | reads) = 0. Then, for a given CpG site *s* in Cell A and Cell B, the respective copy numbers *c*_A,s_, *c*_B,s_, and respective reads, we can compute the expected distance over the possible combinations of allele status between the two cells at *s* (Equation 3):

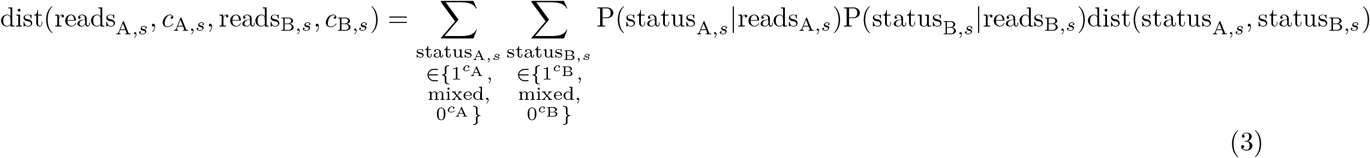

where dist(11,11)=dist(10,10)=dist(00,00)=0, dist(11,10)=dist(10,11)=dist(00,10)=dist(10,00)=0.5, and dist(11,00)=dist(00,11)=1.

The total expected distance between cells A and B can now be computed with some distance function over the vector of expected distances over all shared sites. The *L*_1_ norm normalized by the number of shared sites is computed via Equation 4. The use of shared CpG sites between pairs of single cells to estimate their distances was established by Hui *et al*. [19]:

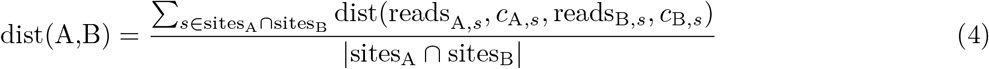

A comparison of our distance formulation against a baseline distance measure commonly used in prior studies [5, 14] on simulated data can be found in Supplementary Section S2.1. After computing the distance between each pair of cells, one can incorporate any distance-based tree reconstruction method into `Sgootr` to obtain the lineage tree and arbitrarily choose a single cell from the normal tissue as the root. We use the `scikit-bio` implementation of the neighbor-joining (NJ) algorithm [35] for our main analysis. It is possible to replace NJ with an alternative distance-based tree construction method in `Sgootr`. We compare `Sgootr`’s performance while using FastME 2.0 [25], a popular alternative, against that using NJ in Supplementary Section S4 and show that they lead to similar results.

### 2.4 Pruning of CpG sites according to a tree-based methylation status persistence measure

The main body of `Sgootr` consists of an iterative procedure: at each iteration, it (i) computes the pairwise distances among single cells to form the tumor lineage tree using a distance-based algorithm, then (ii) measures the methylation persistence score of each CpG site along the tumor lineage tree and prunes away a fraction of CpG sites that have the least scores before continuing onto the next iteration, and (iii) outputs the tumor lineage tree of the particular iteration whose distance from the tumor lineage trees obtained in consecutive iterations are minimum possible.

We have described how to compute pairwise distance among single cells in Section 2.3, and we use the widely used Robinson-Foulds (RF) distance [34, 37] to measure the differences among the tumor lineage trees constructed on the same set of cells in consecutive iterations. In this section, we focus on how we measure the methylation persistence of CpG sites at each iteration given the tumor lineage tree obtained for these CpG sites. In particular, we define the *methylation persistence score* for a CpG site given a tumor lineage tree (Equation 6): the higher the methylation persistence score for a CpG site, the more stably maintained its methylation status change along a tumor lineage tree.

To facilitate the analysis, we assign to each CpG site covered in each cell its most likely methylation status: given sequencing error-corrected reads from component (ii) of `Sgootr`, we call a site homozygously methylated if all of its reads have the site methylated, homozygously unmethylated if all unmethylated, and heterozygous if the read status for the site is mixed. The status remains unknown if there are no reads covering the CpG site. After this step, we measure the persistence of each CpG site independently: first at each branch of the tree, then for the overall tree.

### Methylation persistence score for a CpG site at a particular branch in the lineage tree

Each branch in the methylation tumor lineage tree induces a bipartition of the tree and subsequently of the leaf nodes, which represent single cells. In other words, let *TN* denote the full set of leaf nodes in a methylation tumor lineage tree *t*, cutting a branch *b* in the tree creates disjoint subsets of leaf nodes *TN*_b_ and 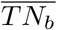. Let *Q*_s,tn_ denote the probability distribution across three possible methylation statuses - homozygously methylated, heterozygously methylated, and homozygously unmethylated - at CpG site *s* across a subset of cells *tn*, and let *I*_s_ represent the set of cells with status information at site *s*. Then, we define *ℳ 𝒫*_*s,b*_, the methylation persistence score of CpG site *s* at branch *b* (Equation 5):

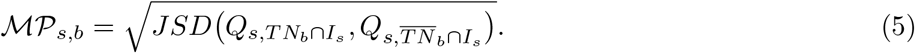

Here *JSD*(*X, Y*) denotes the Jensen-Shannon divergence [39, 26] for a pair of distributions *X, Y*, and is defined as 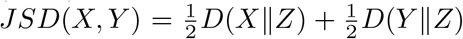 with 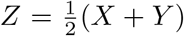 and *D*(*K*∥*L*) denoting the Kullback-Leibler divergence measure for any arbitrary distributions *K* and *L* [22]. It is worth noting that the square root of the Jensen-Shannon divergence measure, which computes *ℳ 𝒫*_*s,b*_ (Equation 5), is metric [32, 12] and is commonly referred to as Jensen-Shannon distance in popular implementations.

The intuition is that, if the change in methylation status of CpG site *s* has an observed persistent effect in tumor progression - namely, *s* is lineage-informative - there exists a branch *b*^∗^ in the methylation tumor lineage tree such that the leaf nodes in the two subtrees induced by *b*^∗^ show very different distributions of methylation statuses. In other words, *ℳ 𝒫*_*s,b*_*∗* will be large for such *b*^∗^. In contrast, suppose the bipartitions contain the same distributions for methylation statuses for *s* - either that most cells share the same status, or that both bipartitions are similarly heterogeneous - the score will be low.

### Overall methylation persistence score for a CpG site in the lineage tree

We define the overall methylation persistence score of CpG site *s* in methylation tumor lineage tree *t* as the maximum methylation persistence score the site *s* has across all valid bipartitions (Equation 6). We recognize that (i) an extreme difference in the number of cells between the bipartitions, or (ii) a severe lack of cells with status information could both lead to meaningless divergence measurements. Therefore, we only consider bipartitions induced by branch *b* such that (i) both partitions contain no fewer than a user-defined fraction *δ* of total number of leaves, and (ii) there is read information in no fewer than a user-defined fraction *ω* of cells in both partitions. In our experiments, we set *δ* = .05 and *ω* = .5.

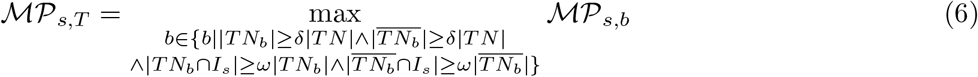

#### Algorithm 2 Partial order-guided modification to the Fitch’s top-down node label assignment

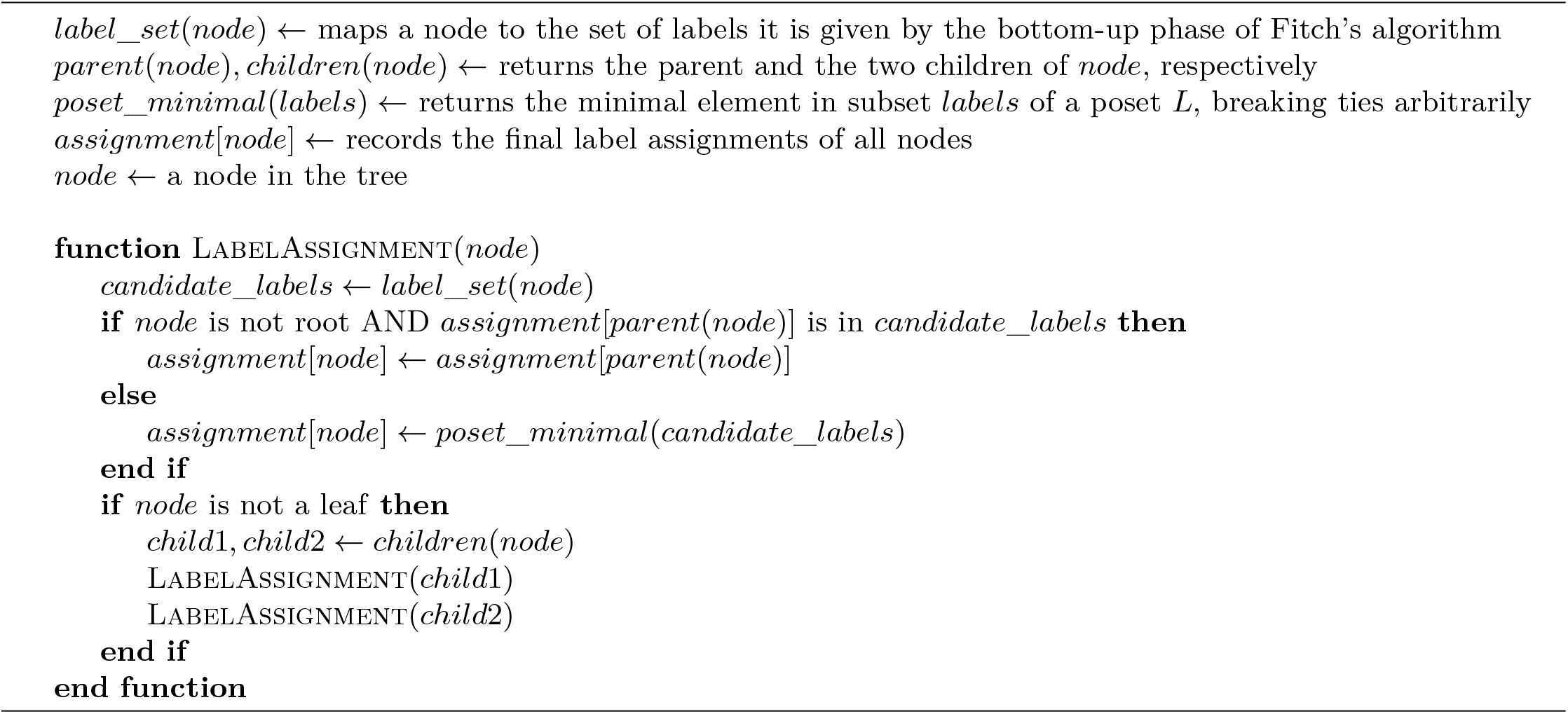

### Iterative joint tumor lineage tree reconstruction and lineage-informative site identification

In a given iteration *k*, `Sgootr` first computes the tumor lineage tree *t*_*k*_ with the distances between pairs of cells (Section 2.3) based on the persistent sites identified in iteration *k* − 1 (for iteration 0 the entire set of sites remaining after component (i) and (ii) of `Sgootr` are used). Then, for each CpG site *s* used in computing *t*_*k*_, `Sgootr` calculates its overall methylation persistence score 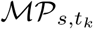. Among the CpG sites with overall methylation persistence scores, `Sgootr` prunes away a user-defined fraction *κ* of the CpG sites with the lowest scores, along with those with tying scores at the threshold. It outputs the remaining CpG sites to be used in iteration *k* + 1. The process continues for a user-defined maximum number of iterations, and `Sgootr` outputs *t*^∗^ = *t*_*i*_ where *i* is the last iteration with *RF* (*t*_*i*_, *t*_*i*_ − _1_) equals the global minimum across iterations.

Intuitively, we would like to detect an approximate point in the iterative procedure where most non-lineage-informative CpG sites have been pruned out (leading to the initial roughly decreasing trend of RF distance) but most lineage-informative CpG sites still remain (whose further elimination will lead to increasingly inaccurate distance measurements between cell pairs and therefore increasingly unstable tree topologies, which is reflected in a once-again increasing trend of RF distance). We demonstrate empirical observations corroborate with such intuition (Figure 1c.,S9,S15) and provide practical recommendations for choosing *κ* and the maximum number of iterations in Supplementary Section S7.

**Fig. 1:**
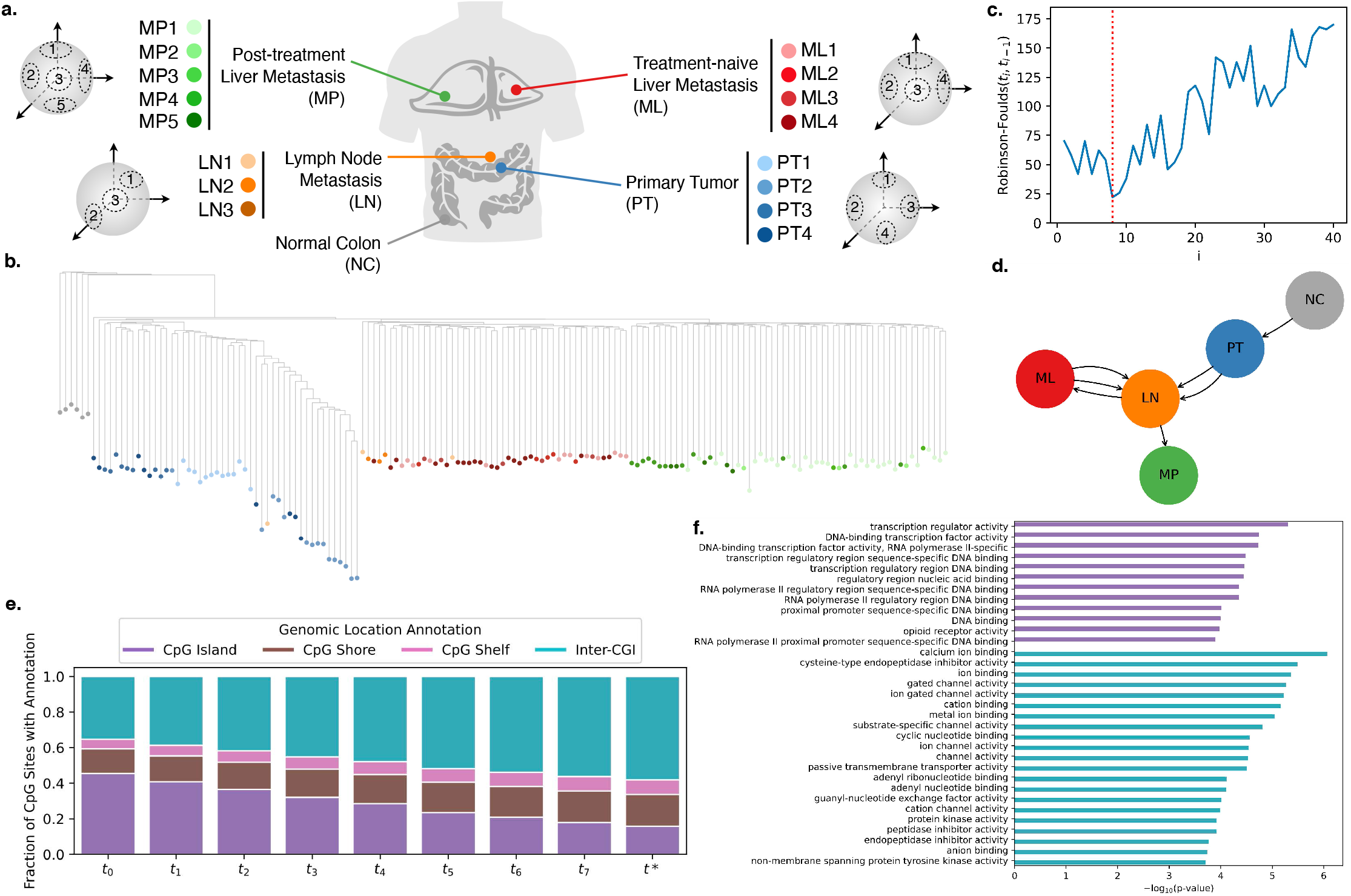
Application of Sgootr to patient CRC01 scTrio-seq data by Bian *et al*. **a**. The multiregional patient data consists of single cells sampled from 4 distinct lesions (in addition to normal tissue) and multiple sampling locations within each lesion. **b**. Tumor lineage tree constructed by `Sgootr` with *κ* = .1. Each single cell is represented as a leaf, colored by its sampling location. **c**. RF distances between the trees of consequtive iterations of the pruning procedure, with global minimum occurring at *t*∗ = *t*_8_. **d**. Tumor migration history inferred by `Sgootr` for patient CRC01. **e**. Fraction of CpG sites located in CpG island, CpG shore, CpG shelf, and inter-CGI regions in the CRC01 trees at each iteration of the pruning procedure of `Sgootr`, from *t*_0_ to *t*∗. **f**. GO terms with significant (*<* .05) q-values in enrichment analysis of non-pseudo, protein-coding genes spanning lineage-informative sites in CpG island and inter-CGI regions respectively in CRC01.

### 2.5 Inference of migration history from lesion-labeled tumor lineage tree

Given a rooted tumor lineage tree *t*^∗^ produced by `Sgootr` through the procedures described above, provided that the single cells in the tree are labeled by their lesion of origin, we can infer the underlying *tumor migration history*, which we represent as a directed multigraph (without self loops) where each vertex represents a distinct lesion and each edge represents a distinct migration event from the source lesion to the target lesion. We accomplish this by first applying the well-known Fitch’s algorithm [13] with slight modification to obtain a unique, maximally parsimonious labeling of the internal nodes of *t*^∗^, then identifying the migration events.

One may recall that, the bottom-up phase of the Fitch’s algorithm computes in a tree the number of migration events - namely, the change of lesion of origin labels from a parent node to a child - in the most parsimonious labeling of its internal nodes. This phase also produces a set of possible labels for each internal node in the tree. The top-down phase, then, assigns each internal node a label from its set, as described in Algorithm 2, starting with root node of *t*^∗^.

The most general formulation of Fitch’s algorithm may produce multiple optimal labelings of the internal nodes; however, some of these optimal labelings may adhere to our prior understanding of the sequential-progression (a.k.a. metastatic-cascade) model of tumor evolution [24] better than others. One can encode such prior understanding as a partial order over the lesion of origin labels of single cells. For example, in this work, we leverage the following partial order:

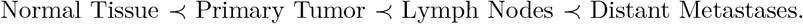

In our modification to the Fitch’s algorithm, during the top-down phase, when a node cannot inherit the label of its parent, we assign it the minimal element in its candidate label set according to the partial order, breaking ties arbitrarily when the elements in the label set cannot be compared (Algorithm 2). This returns a maximally parsimonious labeling of the internal nodes in the tree that also acknowledges the sequential-progression model.

With a fully labeled tree, we produce a directed multigraph that corresponds to the tumor migration history as follows. First, we generate a vertex for each distinct lesion represented in the tree. Then, as we traverse down the tree from the root, for each label change from a parent to a child in the tree, we add one direct edge to the multigraph, from the lesion vertex corresponding to the parent label to that corresponding to the child label. Parallel edges between a pair of vertices correspond to a polyclonal seeding event, and any back edge indicates a reseeding event.

## 3 Results

The performance of `Sgootr`’s distinct components on simulated data are described in the Supplementary Sections S1 (biclustering), S2.1 (distance calculation), and S2.2 (iterative pruning). As can be seen, each of these components substantially improve on the respective baseline approach on a wide range of settings. Furthermore, the results of the iterative pruning component demonstrate that `Sgootr` can capture the migration history of a tumor accurately.

In the remainder of the paper we focus on the application of `Sgootr` to the scBS-seq dataset generated by Bian *et al*. on 9 metastatic CRC patients^8^, among whom 2 (CRC01 and CRC04) have matching scRNA-seq data (via the scTrio-seq2 protocol) from both normal and tumor cells with which copy number calling can be performed [2]. We highlight results from patient CRC01 since in addition to having copy number calls available, they have the largest number of distinct tissue types, sampling locations, and treatment conditions. Full details and results for the other 8 patients can be found in Supplementary Section S3. We additionally present in Supplementary Section S5 `Sgootr`’s results on a GBM patient by Chaligne *et al*.[5], which demonstrate the applicability of our approach to MscRRBS data. To the best of our knowledge, this is the only other single cell bisulfite sequencing dataset with multiregional tumor sampling that is publicly available.

### Sgootr obtains a simple tumor migration history with scBS-seq and scRNA-seq data from patient CRC01

The single cells of CRC01 are sampled from 4 distinct lesions - primary tumor (PT), lymph node metastasis (LN), liver metastasis (ML), and post-treatment liver metastasis (MP) - as well as normal colon tissue (NC) adjacent to the primary lesion; furthermore, there are multiple sampling locations within each lesion (Figure 1a.). We only include cells with both scBS-seq and scRNA-seq data, and employ `inferCNV`^9^ to leverage scRNA-seq data for copy number calling. We apply `Sgootr` with *κ* = .1, terminating the iterative procedure after 40 rounds. A unique global minimum among the RF distances is identified at iteration 8 (Figure 1c.), hence *t*∗ = *t*_8_ (Figure 1b.) is output as the lineage tree. The tumor migration history inferred by `Sgootr` (Figure 1d.) is simpler than previously reported (see Figure S7 by Bian *et al*. [2]): NC grow into PT, followed by a polyclonal migration to LN, which appear to proceed to seed both ML and MP.

While the migration history also suggests a polyclonal reseeding from ML to LN, both edges in the graph have low support: they are both due to a subtree of a particular cellular origin harboring a (potentially mislabeled) singular cell of a different origin.

### Sgootr infers migration histories simpler than alternative methods for Bian *et al*. CRC cohort

We benchmark `Sgootr` against (i) a naïve baseline distance-based tree construction method (column “baseline” in Figure 2a.), (ii) IQ-TREE[30] with the two-state general time reversible model (`GTR2`), an instance of the popular maximum-likelihood-based tool leveraged for inferring lineage trees from single-cell bisulfite sequencing data in prior studies [14, 5] (column “IQ-TREE” in Figure 2a.), and (iii) IQ-TREE (with GTR2 model) preceded by *four-gamete analysis* (FG), a lineage-informative site-selection method previously described by Gaiti *et al*., which returns a subset of input CpG sites which lower than expected epimutation rate [14] (column “FG+IQ-TREE” in Figure 2a.). Note that it is IQ-TREE preceded by FG that was employed exactly in previous studies [14, 5]. However, as discussed in more detail below, the FG step is not only very slow, it also leads to highly complex migration histories. These observations lead us to additionally benchmark against IQ-TREE without the FG step, even though it had not been directly applied to single-cell bisulfite sequencing data in prior studies. See Supplementary Section S8 for further experiment details.

**Fig. 2:**
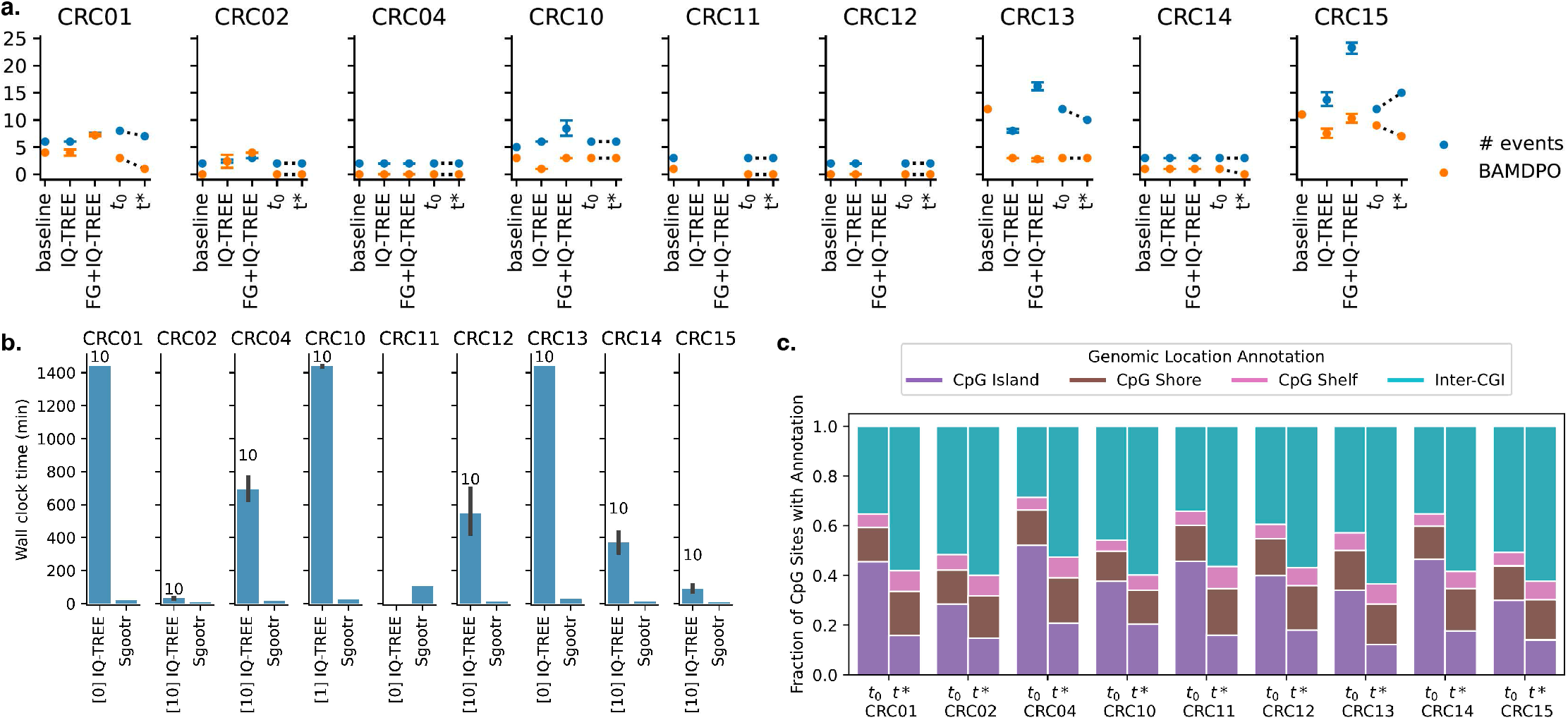
Overview of results from the Bian *et al*. metastatic CRC cohort. **a**. Number of migration events and BAMDPO values of migration histories obtained via (i) naïve baseline method, (ii) IQ-TREE, (iii) IQ-TREE on FG-selected CpG sites, (iv) `Sgootr` intermediate tree prior to iterative procedure (*t*_0_), and (v) final tree output by `Sgootr` (*t*∗). IQ-TREE result is represented by the mean across 10 runs with different random seeds, with the error bars denoting the 95% confidence interval of the mean. The lack of data points in “IQ-TREE” columns means IQ-TREE fail to finish model optimization to start tree search after 24 hours, and that in “FG+IQ-TREE” columns means FG analysis fail to terminate after 100 hours. **b**. Run time comparison between `Sgootr` and IQ-TREE. Each experiment is performed on 4 cores. The number in brackets next to “IQ-TREE” on the x-axis represents the number of IQ-TREE runs (out of 10) that have converged within 24 hours. The number above each IQ-TREE bar represents the total number of runs (out of 10) with any tree search output (converged or intermediate after 24 hours), and hence contributes to the run time and migration history comparison. **c**. Fraction of CpG sites located in CpG island, CpG shore, CpG shelf, and inter-CGI regions, before (*t*_0_) and after (*t*∗) the iterative pruning component of `Sgootr`.

For each benchmarking experiment, we take as input the cell-by-site read count matrices resulting from first-pass heuristic filtering step of `Sgootr`, namely, removing low-quality cells, CpG sites with coverage in less than 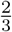 of remaining cells, sites on sex chromosomes, and sites within chromosome 21 peri-centromeric regions. Then, for each CpG site covered in each cell, call whether it is methylated or not by the status receiving support from at least 90% of reads at the site in the cell, discarding entries where such determination cannot be made. This binarization step has been common practice in prior works on single-cell methylation data[2, 14, 5] and is necessary in generating input for the `GTR2` model. For the FG+IQ-TREE experiments, we further perform FG (Supplementary Section S8 for detail) to select CpG sites to input to IQ-TREE. Additionally, we further filter the input to the benchmarking experiments so that all experiments (Figure 2a.) on a particular patient act on the same set of input cells. This calibration is to ensure fair comparisons among results from different methods using the two performance measures we will introduce below. To facilitate fair runtime comparison between IQ-TREE and `Sgootr`, we also filter to ensure the input to IQ-TREE for each patient is the same set as those to `Sgootr`, so that the runtime measurements in Figure 2b. are taken on the same set of cells and CpG sites for each patient.

For each patient in the Bian *et al*. cohort, we generate a baseline lineage tree by first computing the Hamming distance between every pair of cells over their binary CpG site status vectors, averaging across the sites covered in both cells, then applying NJ to the distance matrix. As IQ-TREE is a stochastic method, we perform 10 separate runs of each IQ-TREE experiment with different random seeds, and for each instance report the intermediate tree at convergence or after 24 hours if convergence has yet to be achieved by then.

If none of the 10 runs of IQ-TREE for a particular experiment have finished model optimization, started tree search, and produced some intermediate result, we do not report IQ-TREE result for that experiment. Similarly, if FG does not terminate in 100 hours, we do not report FG+IQ-TREE result for that experiment. We root the baseline and IQ-TREE trees with the same normal cells we used to root the trees by `Sgootr`, and we infer the tumor migration histories from them via `Sgootr`’s modified Fitch’s algorithm.

We compare the migration histories inferred via `Sgootr` and the three alternative approaches by two measures: (i) the total number of migration events, namely the number of edges in our multigraph representation, and (ii) one we name *binary adjacency matrix distance from partial order* (BAMDPO), which is the Hamming distance between the binary adjacency matrix of the simple directed acyclic graph induced by the lesion partial order (see Section 2.5) and that obtained by collapsing all parallel edges in the migration history multigraph. Intuitively, the first measure (output from Fitch’s algorithm) captures the conventional definition of parsimony, and the latter measures the degree the inferred history deviates from the sequential-progression model of tumor evolution. Together, they offer complementary perspectives on the complexity of the inferred histories. For both measures, a lower value would correspond to a simpler history.

Figure 2a compares the tree *t*∗ obtained by `Sgootr` against the alternatives with respect to the two measures. The figure also includes results of *t*_0_ for each patient, the tree obtained by `Sgootr` prior to its iterative site pruning procedure. Comparisons of the measures for *t*_0_ against those for *t*∗ show `Sgootr`’s iterative procedure almost always reduces the complexity of the inferred migration history, provided such a reduction is at all possible. In the case of patient CRC15, even though the total number of migration events increases from *t*_0_ to *t*∗, BAMDPO decreases; this suggests what first appeared to be deviations from the simple sequential-progression model could potentially be resolved as polyclonal seeding events more in concordance with the model, once the tree structure becomes more refined through the iterative procedure. Compared to naïve baseline and the two IQ-TREE-based methods, `Sgootr` typically infers simpler migration histories; the only exception is patient CRC10, where IQ-TREE (without FG site-selection) infers a migration history with the lowest BAMDPO.

### Sgootr-identified lineage-informative CpG sites construct lineages suggesting simpler migration histories than those identified via FG

Specifically, comparisons between the measures from running IQ-TREE on FG-selected sites and those from `Sgootr` show that `Sgootr` consistently outperforms the former in terms of the number of events and BAMDPO where further improvements were possible. This suggests the sites selected by `Sgootr` may be more meaningful for the purpose of lineage reconstruction than those by FG. FG is also time-consuming: for patient CRC11 and CRC12, FG fail to complete after 100 hours, and no experimental results were reported (Figure 2a.).

### Sgootr is orders-of-magnitude faster than IQ-TREE

While `Sgootr` produce as simple, if not simpler migration history than IQ-TREE (without FG site-selection) in terms of the two measures in all but one case, `Sgootr` obtain the results in a fraction of time IQ-TREE took given the same input dimensions. Each IQ-TREE run was allotted 24 hours. If the program converge within 24 hours, we report the wall clock time elapsed; if the program did not converge within 24 hours but have produced some intermediate output from tree search, we report the program run time as 24 hours; and if IQ-TREE fail to produce any intermediate result from tree search within 24 hours, we do not include the elapsed time from that run. Within the allotted time, IQ-TREE produce converged results in 6 out of 9 patients, some (intermediate) result for 8 out of 9 patients, and is not able to finish model optimization and start tree search within 24 hour for any of the 10 runs for CRC11 (Figure 2b.). `Sgootr` is deterministic and obtains lineage trees suggesting simpler histories in a mere fraction of the time when given the same number of cores and memory as IQ-TREE, positioning itself as the better option in more time-frugal settings.

### Sgootr-identified lineage-informative CpG sites are enriched in inter-CGI regions

In all 9 CRC patients in the Bian *et al*. cohort, as well as the multiregional GBM patient MGH105 in the Chaligne *et al*. cohort, the set of lineage-informative sites identified by `Sgootr` showed enrichment in inter-CGI regions (Figures 1e., 2c., S16f.). The median distances between CpG sites and their closest CpG islands also increase as the iterations approach *t*∗ across all patients (Figure S12).

### Genes associated with Sgootr-identified lineage-informative CpG sites are enriched for pancancer and CRC-related terms

To examine whether the lineage-informative sites discovered by `Sgootr` in the Bian *et al*. cohort have potential functional implications, we stratified lineage-informative sites by their genomic annotation with respect to CGI, for each group finding a set of non-pseudo, protein-coding genes whose gene bodies and 1000 bp upstream promoter regions contain the sites, and carried out Gene Ontology (GO)[1, 6] enrichment analyses respectively using GOrilla [11, 10]. All annotated non-pseudo, protein-coding genes in the genome were used as the background set. A hypergeometric test was used to measure the overrepresentation of the genes in associated GO terms, and the Benjamini-Hochberg FDR-adjusted p-values (q-values) are presented. In 5 out of 9 CRC patients, genes mapped to lineage-informative sites within CGI are strongly associated with transcription regulation and DNA binding (Figure 1f., S10,), processes known to be linked to cancer onset[7]. Among informative sites mapped to inter-CGI regions, prominent enrichment in neural activities including ion transport and gated channel activities were observed across all 9 CRC patients (Figure 1f, S10), concurring with recent studies which support the striking cross-talk between the neural system and colorectal cancer development[40, 33]. Enrichment results using an alternative tool, DAVID[36, 18], are also included for comparison (Figure S11). While the GO terms discovered are not identical between the tools, the results highly concur as indicated by the overlap coefficients of the terms (Table S4).

## 4 Discussion

We present `Sgootr`, the first distance-based tool to jointly infer tumor lineage using single-cell methylation data and select for lineage-informative CpG sites. We show `Sgootr` is able to produce simpler tumor migration history, perform CpG site selection, and obtain results orders-of-magnitude faster than alternative methods in applications to real multiregionally sampled single-cell methylation data in CRC and GBM. `Sgootr` concomitantly tackles two key challenges in tumor migration history inference from single-cell methylation data: sparsity and diverse inheritance dynamics of CpG methylation. `Sgootr` provides a fast and reliable solution to leverage single-cell methylation profiles and elucidate the complex tumor epigenetic landscape.

On top of tumor lineage inference, `Sgootr` captures meaningful biological insights of lineage-informative CpG sites. In both cancer types included in our study, CRC and GBM, we discovered the enrichment of lineage-informative sites in inter-CGI regions across different patients. Interestingly, we also observed that the variability of the methylation status of a CpG site among all sequenced cells seems to increase with its distance from the closet CpG island (Table S5). While the underlying mechanisms of these phenomena are yet to be elucidated and beyond the scope of this study, a plausible explanation is that CpG sites in islands are protected by the reduced chromatin accessibility in the region marked by histone modifications[4], whereas inter-CGI sites harbors more stochastic changes due to malfunctioning of DNA methyltransferases and TET enzymes [9]. While CpG islands have attracted much attention in the studies of tumor epigenetic changes and more frequently covered in methylation arrays, our findings suggest that inter-CGI regions should not be overlooked especially in the context of tumor lineage tracing. We are looking to explore the prognostic potential of inter-CGI CpG sites in our subsequent studies.

## Supplementary Material

## S1 Biclustering: benchmarking on simulated data and a case study on CRC01

As can be seen in Figure S2 (left plot) on the colorectal cancer patient CRC01 [2], the distribution of the number of sites covered by at least one read across cells is bimodal and a relatively small fraction of cells have very few sites covered. In contrast, the distribution of the number of cells with at least one read covering a site (right plot) is unimodal across the sites and only a relatively small fraction of sites are covered in a large number of cells. Provided the presence of those well-covered sites are distributed over the cells independently and identically (i.e. in i.i.d. fashion), the heuristic filtering described in Section 2.1 would work well. However, if the distribution is not i.i.d., fixing the output size, the submatrix obtained by ILP-based biclustering formulation would have higher density, improving the overall reliability of the tree constructed by `Sgootr`. Since, for given *α* and *β* values, the ILP formulation can only produce a submatrix no sparser than that produced by the heuristic filters, it is our preferred choice. Below we explore how the running time and memory usage of the ILP formulation vary with key problem parameters, and identify under what parameter settings of the problem instance it remains feasible.

### S1.1 Running time and memory benchmark on simulated data

We first demonstrate how the expected running time (in other words, time the Gurobi solver takes to obtain an optimal solution) and memory behaviors of the Gurobi solver on our formulation vary as a function of the input size.

For the problem at hand, the input size is parameterized by *v*, the number of cells, and *u*, the number of CpG sites in the input matrix *M*. The number of variables (*N*_var_), the number of constraints (*N*_constr_), and the number of non-zero entries in the constraint matrix (*N*_nz_) are some features of the Gurobi representation of the linear program that may influence its memory usage and running time; note that all three features are *𝒪* (*vu*) (Table S1). Also note that Gurobi uses the sparse representation for the constraint matrix; as a result, even though the naive representation of the constraint matrix requires *𝒪* (*v*^2^*u*^2^) space, the size of its representation by Gurobi is proportional to *N*_nz_, which is *𝒪* (*vu*).

Below we proceed to simulate datasets with varying sizes and study their running time and memory usage.

**Table S1:** Feature sizes of the biclustering formulation model as a function of the number of cells, *v*, and the number of CpG sites, *u* in the input.

| Feature | $f(v, u)$ | Asymptotics |
| --- | --- | --- |
| # Variables ($N_{var}$) | $v \times u + v + u$ | $\mathcal{O}(vu)$ |
| # Constraints ($N_{constr}$) | $3 \times v \times u + 2$ | $\mathcal{O}(vu)$ |
| Size of constraint matrix | $N_{var} \times N_{constr}$ | $\mathcal{O}(v^2u^2)$ |
| # Non-zeros ($N_{nz}$) | $7 \times v \times u + v + u$ | $\mathcal{O}(vu)$ |
| Density of constraint matrix | $N_{nz} / (N_{var} \times N_{constr})$ | $\mathcal{O}((vu)^{-1})$ |

### Simulation details

For simplicity, we set *v* = *u* in our simulations. For each (square) input binary matrix *M*_sim_, we set a “dense” submatrix of size .7*v* ×.7*u* such that each entry in the submatrix is set to 1 independently with probability *q*, while each entry outside of the submatrix is set to 1 with probability 1− *q*. (An entry set to 1 indicates that the corresponding cell has at least one read covering the corresponding site.) We vary the simulation parameters as follows:

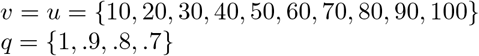

For *q* = 1, we have a .7*v* ×.7*u* submatrix with all 1’s and 0’s everywhere else. As *q >* .5 decreases, while we still have a dense .7*v* ×.7*u* submatrix, its density difference with the rest of the matrix is not as pronounced, making it more difficult to achieve our objective. With each set of parameters (*v, u, q*), 5 matrices are simulated (for investigating the stochasticity introduced by *q <* 1), resulting in a total of 10 × 4 × 5 = 200 simulated matrices.

## Results

We perform biclustering on each of the 200 simulated binary matrices with *α* = *β* ={.6, .7, .8}, resulting in 600 trials total. (We remind the reader that *α* and *β* are respectively the proportion of the rows and columns of the densest submatrix we aim to identify.) The trials are performed serially on the same machine with a single thread. As the dense submatrix in each simulated matrix has dimensions .7*v* × .7*u*, performing biclustering with *α* = *β* = .6 and *α* = *β* = .8 will respectively correspond to an underestimation and overestimation of the two parameters. We profile (i) running time and (ii) peak memory usage for all trials.

As a general pattern, both wall-clock running time (Figure S1a., where the y-axis is log_2_-transformed to better visualize the differences) and peak memory usage (Figure S1b.) grow linearly with *v* × *u*. For any given input size, the running time increases with decreasing value of *q*, as the objective becomes more difficult to achieve (as demonstrated within each column in Figure S1a.). Notably, the running time also seems to be impacted by the choice of *α* and *β*: while both underestimation and overestimation lead to an increase of running time possibly due to the existence of several equally good solutions, underestimation seems to lead to a more substantial penalty (as can be observed across columns corresponding to the same input size in Figure S1a.). The peak memory usage seems to be similar for a given size input across choices of *α* and *β*, though an underestimation seems to occasionally lead to considerably higher usage and hence higher variance across trials (as can be observed across columns corresponding to the same input size in Figure S1b.).

**Fig. S1:**
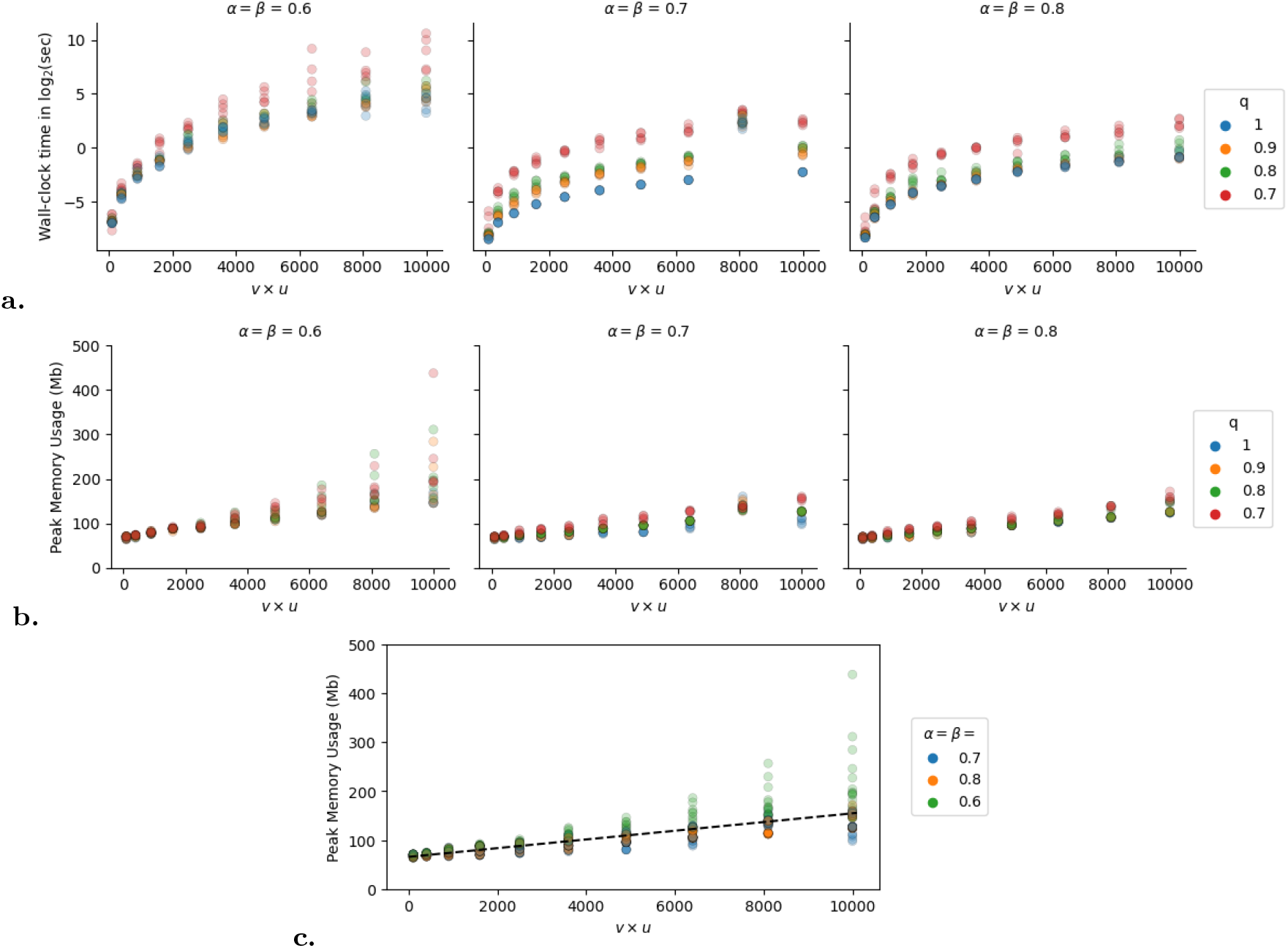
Running time and memory behavior on 200 simulated square binary matrices. **(a.)** Wall-clock time in log_2_ of seconds. Measures were log_2_-transformed to better visualize the difference. **(b.)** Peak Memory consumption in Mb. **(c.)** Linear regression on peak memory usage across all 600 trials. Dashed line describes *y* = .0089*x* + 65.69.

To allow extrapolation of peak memory usage for any given input size, we perform linear regression using the ordinary least squares method on the data from the 600 trials, with *v* × *u* as the independent variable and peak memory usage in Mb as the dependent variable, obtaining a *y*-intercept of 65.59 and slope of .0089.

We do not extrapolate running time in the same manner, as it varies drastically with problem difficulty, which we proxy with *q* in our simulation study, but is hard to estimate in real data. Running time will also depend on the specific optimization strategies employed by Gurobi, a proprietary software, that may depend on other features of the problem instance, such as the density of the constraint matrix. Furthermore, one can always specify a time limit for a Gurobi run and obtain an intermediate output and its associated optimality gap, meanwhile Gurobi fails without output if memory limit is reached; therefore, memory is more likely to present a bottleneck for large-scale problems than time in practice.

Results from simulated data lead us to two key conclusions: (i) memory usage may be a critical limiting factor to the feasibility on large input, and (ii) well-chosen *α* and *β* values will not only yield a more informative submatrix for the remainder of the `Sgootr` workflow, they will also (typically) lead to faster converge than ill-chosen parameters.

### S1.2 Biclustering case study on CRC01 from the Bian cohort

Next, we investigate the performance of our biclustering formulation on the CRC01 dataset from the Bian cohort as we vary *α* and *β* values.

### Data descriptions

The dataset has in total 409 single cells with scBS-seq data from patient CRC01, among which 241 also has matching scRNA-seq data. As we will call copy number from scRNA-seq data to input to `Sgootr`, we only include those 241 cells with both scRNA-seq and scBS-seq data for the biclustering procedure. Collectively, the 241 cells cover 52,343,774 unique autosomal CpG sites. Figure S2 shows per-cell CpG site coverage distribution and per-site cell coverage distribution. The induced 241 × 52, 343, 774 binary coverage matrix has a density of 13.21%.

**Fig. S2:**
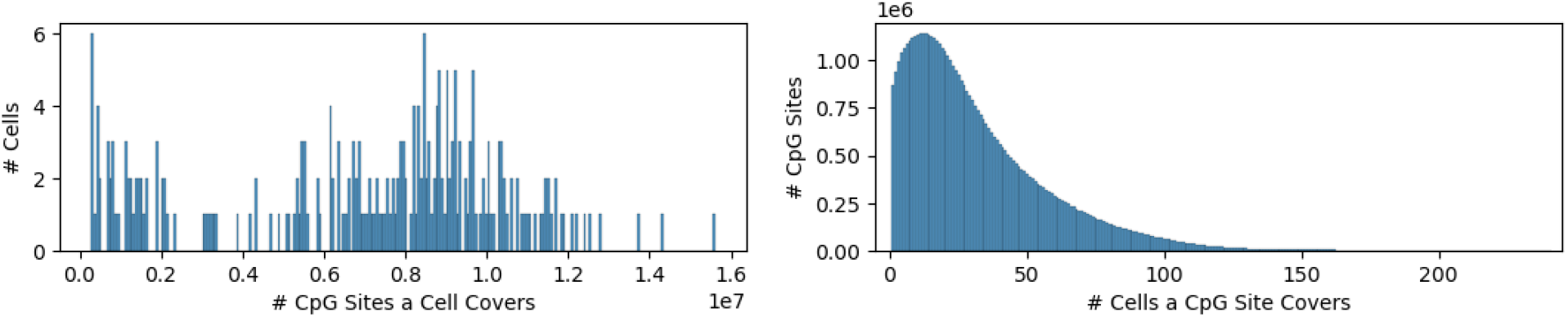
Patient CRC01 cell (left) and site (right) coverage level distribution.

**Table S2:** Densities of the original binary matrix and the subsampled matrices.

| Original | Subsamples (24 × 52,343) |  |  |  |  |  |  |  |  |  |  |  |
| --- | --- | --- | --- | --- | --- | --- | --- | --- | --- | --- | --- | --- |
| (241 × 52,343,774) | 0 | 1 | 2 | 3 | 4 | 5 | 6 | 7 | 8 | 9 | mean | std |
| **13.21%** | 14.53% | 12.22% | 14.58% | 11.90% | 12.17% | 12.74% | 12.09% | 15.11% | 16.31% | 10.34% | **13.20%** | **1.74%** |

We aim for the resulting submatrix to have a density of at least 66%. The intuition here is that `Sgootr` estimates the distance between any pair of cells using the set of CpG sites which are covered by at least one read in each cell; provided 2*/*3 of the picked sites are covered by a read in each cell, the distance between any pair of cells will be based on at least 1*/*3 of the picked sites, i.e. at least half of the sites covered by a read in each cell. This ensures that the number of CpG sites employed in distance estimations involving one particular cell do not vary by more than a factor of 2. While we do not ensure that each row (or for that matter each column) has a density of at least 2*/*3, an overall density of 2*/*3 (or ∼ 66%) in the submatrix identified by our ILP formulation should provide a reasonable distance estimate between cell pairs.

### Estimating *α* and *β* values

To estimate *α* and *β*, we generate 10 disjoint subsamples of the dataset, each containing 10% of the 241 cells and 0.1% of the 52,343,774 CpG sites. For each subsample, we generate the corresponding 24 × 52, 343 binary matrix (Table S2), on which we perform biclustering with all possible combinations of *α* = {.4, .5, .6, .7, .8, .9} and *β* = {.001, .005, .01, .05, .1, .15} in parallel, resulting in 10 × 6 × 6 = 360 runs of biclustering in total. The choices for the ranges of *α* and *β* values are informed by the density of the original matrix, the desired density of roughly 66%, and the cell and site coverage distributions (Figure S2). We will determine *α* and *β* based on i) time-to-convergence, ii) optimality gap at time limit, and iii) density of resulting submatrix in the subsampled trials.

**Fig. S3:**
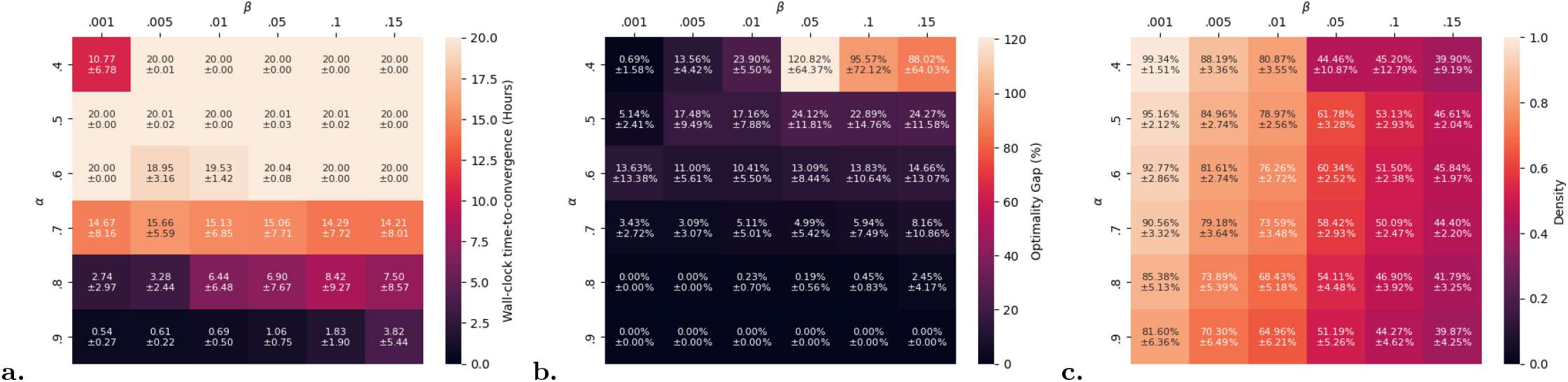
Mean and standard deviation of (a.) wall clock time-to-convergence in hours, (b.) optimality gap at 20 hours, and (c.) resulting submatrix density across the 10 subsamples, over a range of (*α, β*) value pairs. Note that we set the time limit for the biclustering program to 20 hours, and therefore wall clock time-to-convergence is at most 20 hours.

## Results

A total of 360 runs of biclustering were performed across the 10 subsamples over a range of *α* and *β* values. Each run was given 4 threads and 20 hours. Runs with *α* = {.8, .9} were able to achieve negligible optimality gap in a matter of several hours, and smaller values of *α* leads to substantially higher time-to-convergence and optimality gap within the allotted time (Figure S1a.,b.). Fixing *α* = {.8, .9}, looking at the other parameter, runs with *β* = {.001, .005} all solved to convergence with no optimality gap (Figure S1a.,b.). With *α* = {.8, .9} and *β* = {.001, .005}, the resulting submatrices have all achieved a density within a neighborhood of 66%. We choose *α* = .8 and *β* = .005 to i) avoid over-pruning, and ii) considering variance in the different subsamples, have some confidence that results from the full original binary matrix will be at least 66% dense. (Figure S1c.).

The next step would be running biclustering on the original 241 × 52, 343, 774 binary matrix with the chosen parameters *α* = .8 and *β* = .005. However, for input of this scale, the peak memory usage is predicted to be 112 Tb according to our linear regression result from simulated data, and is not feasible on the resources available to us and likely most researchers. Therefore, we opted for performing heuristic filtering in generating the results in the main text. Note that the submatrix we obtained with heuristic filtering on CRC01 is of a similar scale as that from performing biclustering on a 241 × 52, 343, 774 matrix with *α* = .8 and *β* = .005.

For future iterations of the tool, we aim to optimize the implementation of the biclustering procedure to improve memory efficiency, and a combination of heuristic filtering and biclustering may be needed for best solving the problem at hand. In particular we aim to utilize the ILP formulation on the entire set of cells but only a subset of sites so as to stay within the limits of Gurobi’s ability to solve the problem. Given the subset of cells identified by the ILP formulation, we will then be able to employ the heuristic filtering to identify the subset of sites with high coverage on these cells.

## S2 Benchmarking Sgootr on simulated data

In this section we evaluate the performance of distinct components of `Sgootr` on simulated data, in comparison to baseline approaches. All `Sgootr`-related parameter settings will be the default configuration. Furthermore, for simplicity, we will assume copy number = 2 in our simulations.

### S2.1 Expected distance calculation on simulated data

First we focus on our formulation for the expected distances between cell pairs and evaluate its performance against a simple baseline approach that was utilized in earlier studies [14, 5] which ignores sites that are potentially heterozygously methylated.

### Simulation details

Our simulations generate pairs of diploid cells with length-normalized ground truth distance of *π* = {.1, .2, .3, .4, .5, .6, .7, .8, .9}. For each value of *π* we simulate 50 cell pairs, producing in total 9 × 50 = 450 simulated cell pairs. Each cell starts with 2 (CpG) alleles of 1000 unmethylated CpG sites. For any simulated pair, given *π*, we uniformly sample the number of sites where one cell is homozygously methylated but the other is homozygously unmethylated (each contributing 1 to the total distance) and the number of sites where one cell is homozygous but the other is heterozygously methylated (contributing .5 to the total distance) from all possible combinations. For example, with *π* = .6, by pigeonhole principle we have at least 200 and at most 600 CpG sites each of which contributing 1 to the total distance. After sampling those sites, the remainder of the ground truth distance need to be contributed by sites where one cell is heterozygous but the other is homozygous. After sampling this second set of sites, the ground truth distance is already achieved and the remaining sites will be of the same methylation status across the two cells.

Next, for each of the 450 cell pairs with known normalized ground truth distances, we simulate methylated and un-methylated read count vectors for either cell (i) for a range of possible depth of coverage to study how that may impact the measurements, and (ii) using the empirical per-site read coverage distributions from patient CRC01, the key patient we analyze in the main text. For (i), specifically, we model the number of reads covering a CpG site by a zero-truncated Poisson distribution (ZTPois) parameterized by *λ* = {1, 2, 3, 4, 5, 6} (Figure S4a.). For (ii), we obtain the empirical per-site read coverage from patient CRC01 after first-pass filtering (Figure S4b.). The empirical distribution is truncated at 20, as probability substantially diminishes for larger values. For each simulated cell pair, we simulate 5 sets of methylated and unmethylated read count vectors for every depth of coverage distribution, obtaining 450 × 7 × 5 = 15, 750 sets of read count vectors.

We compare the distances between the read count vectors of a cell pair obtained by (i) a baseline distance measure, which we will describe in detail in the following section, and (ii) our expected distance formulation with default parameters. This comparison will help us better understand examine (a) how the depth of coverage affects the two distance measures, and (b) given the empirical read coverage distribution, which measure better models the ground truth distance.

## Results

The baseline distance measure we leverage in the comparison is one commonly used in prior studies [14, 5]. For each CpG site in a single cell, this approach first calls its status as methylated or unmethylated depending on which has with support from at least 90% of the observed reads at that site for the cell; it then discards all sites for which such binarization cannot be done. As a result, each cell will have a binary vector (likely with some missing entries due to failed binarization) indicating whether each site is methylated. Then, for each pair of cells, this approach computes the Hamming distance between the respective binary vectors over all sites at which both have a binarized status, then normalizes the distance by the number of such sites. This distance measure essentially excludes heterozygosity from consideration.

**Fig. S4:**
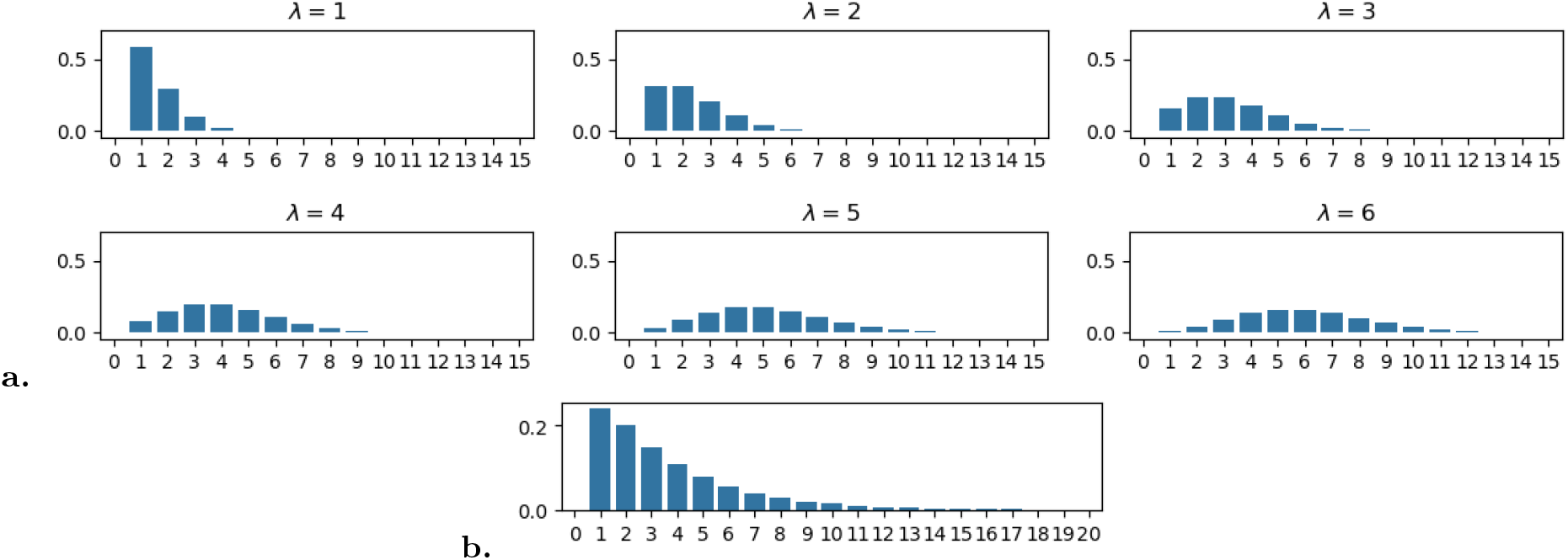
Probability mass functions of (a.) ZTPois(*λ*), where *λ* = {1, 2, 3, 4, 5, 6} and (b.) the empirical per-site read coverage distribution from patient CRC01.

**Fig. S5:**
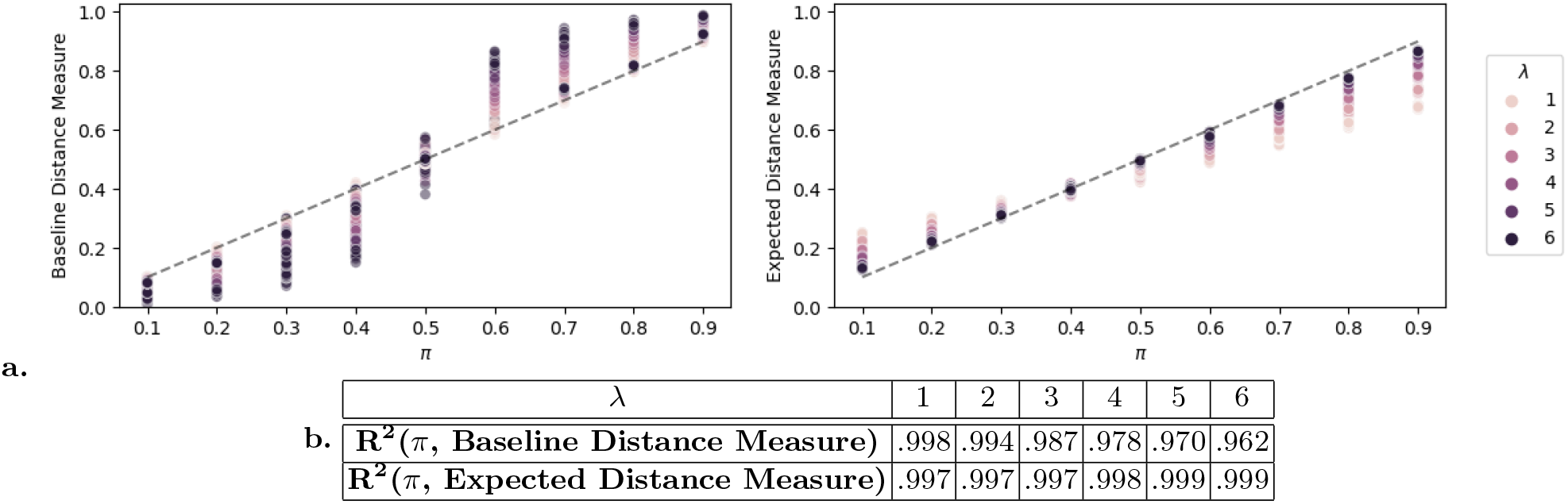
Baseline distance measure and expected distance measure on 13, 500 sets of read count vectors, simulated from cell pairs with known normalized ground truth distance *π* = {.1, .2, .3, .4, .5, .6, .7, .8, .9 }, with level of read coverage depth modeled by ZTPois(*λ*), where *λ* = {1, 2, 3, 4, 5, 6 }. **(a.)** Measured distances across increasing value of *π* for various *λ*. The dashed lines describe *y* = *x*, and are drawn to highlight the direction of deviation in the measured distances. **(b.)** For a given level of read coverage depth parameterized by *λ*, the *R*^2^ values of linear regressions with the base line v.s. our expected distance measures as the independent variable and the true distance *π* as the dependent variable.

**Fig. S6:**
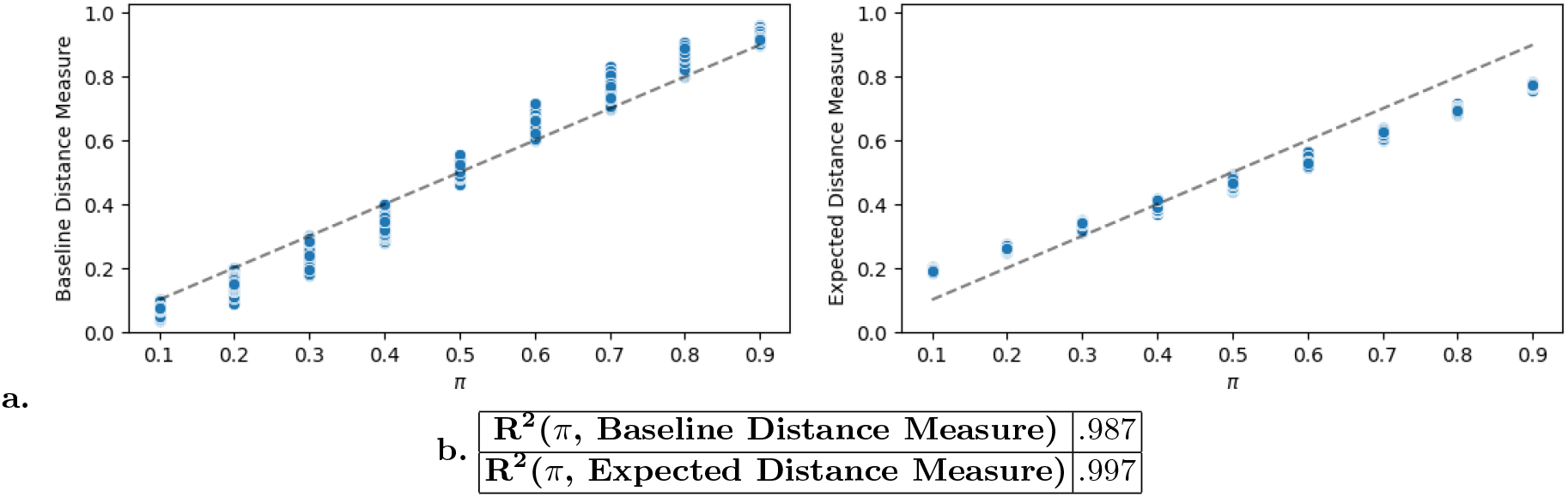
Baseline distance measure and expected distance measure on 2, 250 sets of read count vectors, simulated from cell pairs with known normalized ground truth distance *π* = {.1, .2, .3, .4, .5, .6, .7, .8, .9}, with level of read coverage depth modeled by the empirical read coverage distribution gathered from the CRC01 sample. **(a.)** Measured distances across increasing value of *π*. The dashed lines describe *y* = *x*, and are drawn to highlight the direction of deviation in the measured distances. (Note that the expected distance measure is based on the default configuration with the probability of a site being heterozygous set to 1*/*3.)**(b.)** *R*^2^ values of linear regressions with the measured distances being independent variable and the true distance *π* being the dependent variable.

We first examine how the depth of coverage (whose distribution is as per Figure S4 panel a) affects the baseline in comparison to our expected distance measure. For each of the 450 × 6 × 5 = 13, 500 sets of read count vectors sampled according to ZTPois(*λ*), we measure both distances. The baseline distance measure tend to underestimate the ground truth distance when *π <* .5 and overestimate it when *π >* .5 (Figure S5a., left). This is because 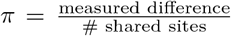 and the exclusion of each unbinarized (i.e. observed to be heterozygous) site subtracts .5 from the numerator, and subtracts 1 from the denominator. This directional deviation from true *π* becomes more substantial as read depth increases, witnessed by the decreasing *R*^2^ values resulting from performing linear regression of *π* on the measured baseline distance (Figure S5b., top row). Intuitively, for small *λ*, likely only one allele at the heterozygous site will be sampled, and (assuming there is no preferential sampling of alleles of a particular status) its status will with 50% chance be called methylated and 50% unmethylated, and on expectation, a heterozygous status and a homozygous status will still contribute an expected distance of .5; however, for larger *λ*, heterozygosity is more likely to be reflected in the observed reads, therefore a larger portion of true heterozygous sites will be excluded, leading to a larger deviation of measured distance from *π*.

In contrast, for any given *λ*, our expected distance measurement has persistently high *R*^2^ values in a linear regression of *π* (Figure S5a., right; S5b., bottom row). In fact, the *R*^2^ values slightly improve with increasing level of coverage. We acknowledge that the expected distance measure, too, deviates from the ground truth *π*, especially at low coverage. As described by the intuition previously provided in Section 2.3, with few reads at a CpG site, it is unlikely (quantified by the user-defined parameters) that they have sampled from all alleles present; therefore, if a CpG site in a cell only has reads of one methylation status, and the number of reads is low, our expected distance measure will assign some probability to it being heterozygous. This will lead to overestimation of the distance at a CpG site if two cells have the same homozygous status but are low-coverage for the site; similarly, this will lead to underestimation of the distance at a CpG site if two cells have the opposite homozygous statuses but are low-coverage for the site. There are proportionally more of the former type of sites for lower values of *π* and more of the latter type of sites for higher values of *π*, explaining the direction of deviation from *y* = *x* (Figure S5a., right). That said, as the coverage level increases, the expected distance measure converges to the ground truth distance *π*.

Since it is reasonable to assume that the per-site read count distribution across all single cells sequenced with the same technology is close to being identical, our expected distance formulation is preferable to the baseline measure, as it tracks the true distance well for any given read coverage level. This is the case especially for higher coverage levels increases, which should be desirable if not destined as sequencing technology improves.

Next, we examine the performance of the two distance measures on data simulated using the empirical read coverage distribution (as per Figure S4 panel b). For each of the 450 × 1 × 5 = 2, 250 sets of read count vectors, we again measure distance using (i) the baseline measure and (ii) our expected distance measure. The trends previously observed with read counts sampled from ZTPois(*λ*) still stand (Figure S6a.), and our expected distance measure again tracks the ground truth distance *π* better at this level of coverage, resulting a higher *R*^2^ value (Figure S6b.).

### S2.2 Iterative pruning on simulated data

Next, we evaluate our iterative pruning approach on simulated data against a baseline approach as described in Section 3.

### Simulation details

We simulate 10 clone trees with different topologies, various degrees of branching, and various depths (Figure S7a.). Each clone tree has 1 normal clone (grey “ROOT” nodes in Figure S7a.) and 20 tumor clones (numbered nodes in Figure S7a.). Each clone is represented by a pair of ground truth lineage-informative CpG-alleles with 9,000 CpG sites. 3000 of these are infinite-sites-abiding, another 3,000 “invariant”, and the final 3,000 those of the “mixture” model [28]. For each clone tree, we first focus on the 3,000 infinite-sites-abiding CpG sites. At “ROOT”, all of these sites are set to be homozygously unmethylated, and at each branch in the clone tree, a subset of these CpG sites that have not been previously selected are selected to have their statuses change from homozygously unmethylated to (i) homozygously methylated with 50% chance and (ii) heterozygously methylated with 50% chance. The subsets are roughly equal in size. The remaining 6,000 CpG sites are simulated at the single cell level, as described below.

Given the clone tree with known (sub)clonal methylation status changes for the “infinite-sites-abiding” CpG sites, we simulate single-cell datasets with added noise and sparsity. To simulate one single-cell dataset from a clone tree, we simulate 10 diploid cells from each clone (21 × 10 = 210 cells for each dataset), each with 6,000 additional CpG sites - the methylation status of the 3,000 “infinite-sites-abiding” sites are copied from their clone of origin. Among the additional CpG sites, the first 3,000 are “invariant” whose methylation status in each cell is with 80% chance homozygous (half of these homozygous sites are methylated by default and the other half are unmethylated), 10% chance heterozygous, and 10% chance the alternative homozygous. The remaining 3,000 sites are of the mixture model of methylation dynamics[28]: each is with 50% chance heterozygous, 25% chance homozygously methylated and 25% homozygously unmethylated.

Given such a set of 210 pairs of CpG-alleles, we sample the corresponding 210 × 9, 000 methylated and unmethylated read count matrices according to the empirical read coverage distribution (Figure S4b.) as previously described in Section S2.1. Furthermore, we randomly mask out the same 33% of the coordinates in either 210 × 9, 000 matrix, introducing sparsity observed in real datasets while mirroring the desired input matrix density (previously justified in Section S1.2).

We simulate 5 such single-cell datasets for each clone tree, resulting in 10 × 5 = 50 total pairs of (un)methylated read count matrices that are 66% dense. We leverage those as input to our simulation experiments.

## Results

We compare the performance of `Sgootr` (no biclustering or error correction, *κ* = .1, for a maximum of 20 iterations) on simulated datasets against a baseline approach as described in Section 3 with respect to the trees constructed. Briefly, given the read count matrices, the baseline approach computes the Hamming distance between the binarized methylation status vectors of each cell pair, normalize by the number of sites covered in both cells, and applies NJ on the pairwise distance matrix to obtain a tree. We root the `Sgootr`- and baseline-generated trees with the same set of arbitrarily picked single-cell from the normal clone, and infer clonal evolution histories from the constructed trees using our Fitch’s algorithm-based approach (Section 2.5) with the ground truth topologies (Figure S7a.) as the guiding partial orders. We use the same two evaluation metrics as in Section 3: (i) number of methylation events, and (ii) BAMDPO, which measures the difference between the inferred and the ground truth topologies.

In terms of the two performance measures, `Sgootr` outperforms the baseline approach in datasets generated from 9 out of 10 simulated clone trees, and performs comparably in datasets generated from simulated clone tree 4 (Figure S7b., comparing between “baseline” columns and “*t*∗ “ columns). Furthermore, despite that `Sgootr` many times already has superior performance compared to baseline prior to its iterative procedure, highlighting the benefits of using the expected distance formulation as opposed to the baseline distance measure (Figure S7b., comparing between “baseline” columns and “*t*_0_” columns), the iterative procedure of `Sgootr` seems to lead to additional boosts in performance by refining the signal (Figure S7b., comparing between “*t*_0_” columns and “*t*∗” columns).

**Fig. S7:**
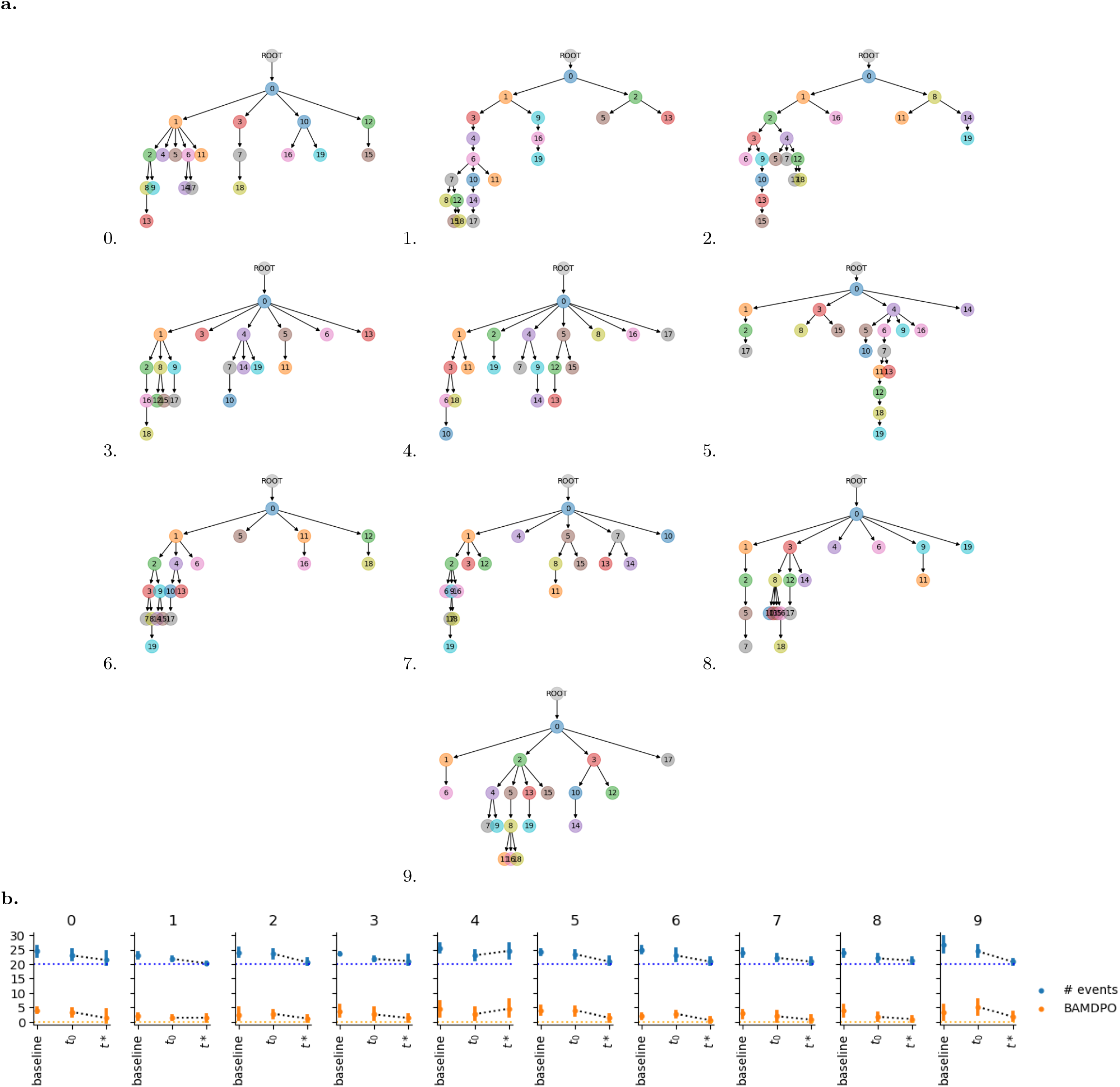
Results on 10 simulated clone trees from Sgootr and baseline method. **(a.)** Topologies of the simulated clone trees. Each clone tree has 1 normal clone and 20 tumor (sub)clones. A total of 3,000 CpG sites abiding by the infinite-sites assumption - and therefore may be lineage-informative - were simulated for each tree. **(b.)** Number of migration events and BAMDPO values of migration histories obtained via (i) naïve baseline method and (ii) `Sgootr` intermediate tree prior to iterative procedure (*t*_0_), and (iii) final tree output by `Sgootr` (*t*∗). Results are represented by the mean across 5 single-cell methylation datasets generated from each clone tree, with the error bars denoting the 95% confidence interval of the mean. Dotted lines in blue and orange mark the ground truth (best possible) values for number of events and BAMDPO, respectively.

## S3 Application of Sgootr on the complete set of 9 metastatic CRC patients made available by Bian *et al*.[2]

**Table S3:** Data summary of the Bian *et al*. metastatic CRC cohort [2]. The number of cells and number of sites are those obtained after component (i) and (ii) of `Sgootr` before entering the iterative procedure.

| Patient | Primary site | Sampled lesions | Number of cells | Number of sites | CNA calls available |
| --- | --- | --- | --- | --- | --- |
| CRC01 | Left Colon | NC, PT, LN, ML, MP | 160 | 140,801 | ✓ |
| CRC02 | Right Colon | NC, PT, ML | 39 | 49,633 |  |
| CRC04 | Right Colon | NC, PT, LN | 65 | 303,849 | ✓ |
| CRC10 | Left Colon | NC, PT, LN | 114 | 275,111 |  |
| CRC11 | Left Colon | NC, PT, LN | 228 | 579,272 |  |
| CRC12 | Rectum | NC, PT, LN | 22 | 650,140 |  |
| CRC13 | Left Colon | NC, PT, LN | 185 | 204,153 |  |
| CRC14 | Rectum | NC, PT, LN | 29 | 511,588 |  |
| CRC15 | Right Colon | NC, PT, LN, ML, MO | 50 | 42,824 |  |

**Fig. S8:**
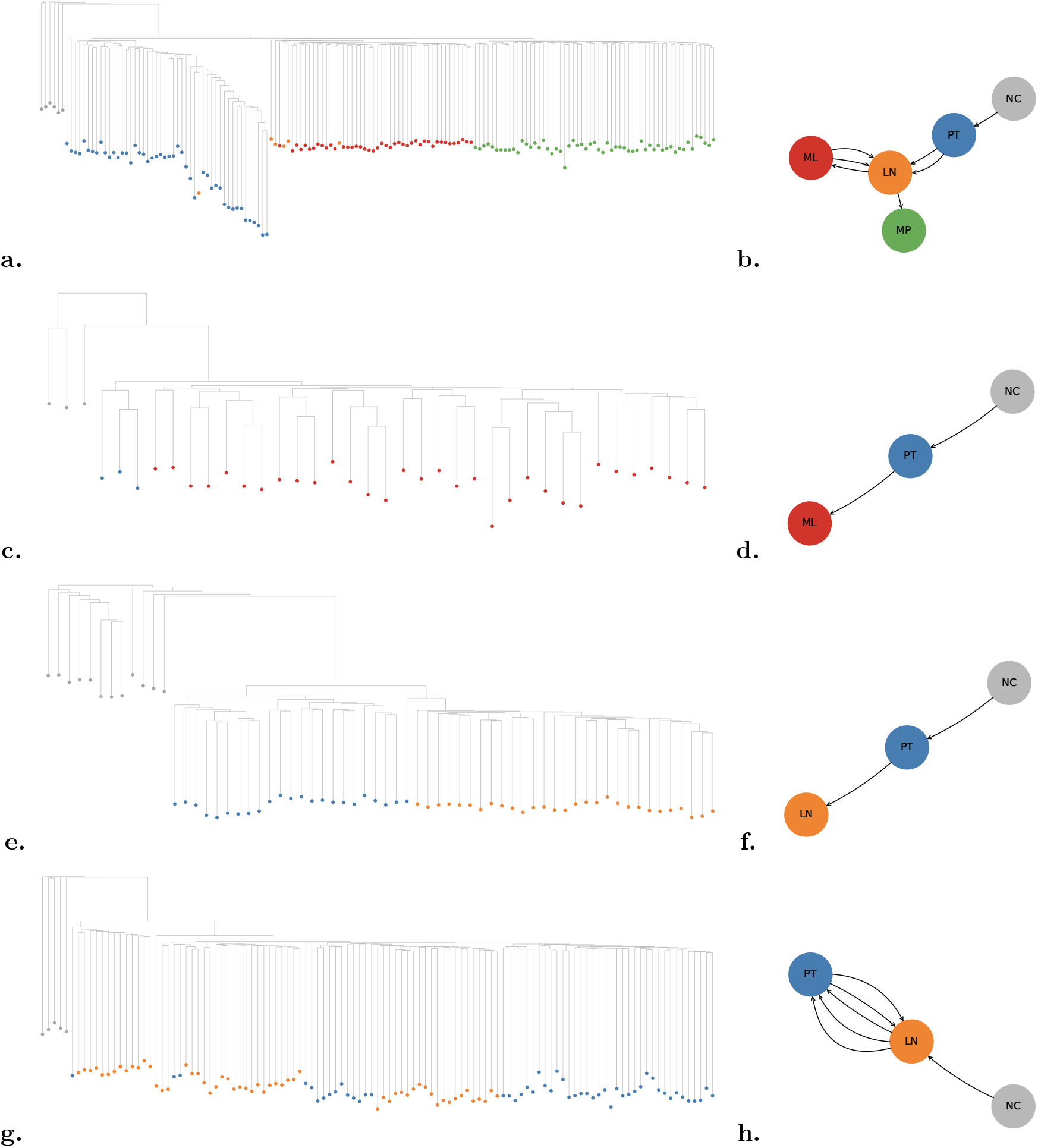
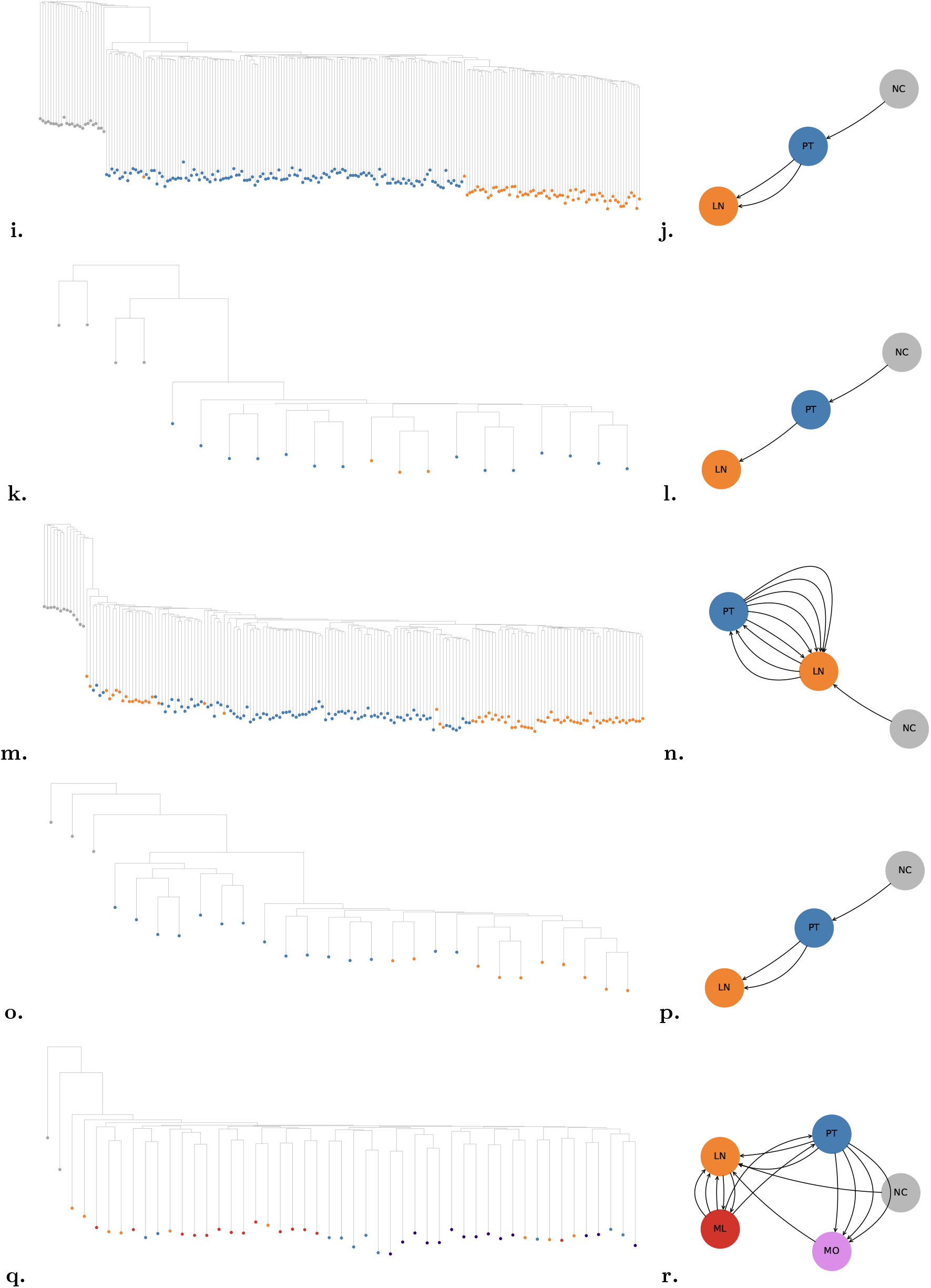
Sgootr-constructed (*κ* = .1) tumor lineage trees (*t*^∗^s) and -inferred tumor migration histories for patients in the Bian *et al*. cohort: CRC01 (**a**., **b**.), CRC02 (**c**., **d**.), CRC04 (**e**., **f**.), CRC10 (**g**., **h**.), CRC11 (**i**., **j**.), CRC12 (**k**., **l**.), CRC13 (**m**., **n**.), CRC14 (**o**., **p**.), CRC15 (**q**., **r**.)

**Fig. S9:**
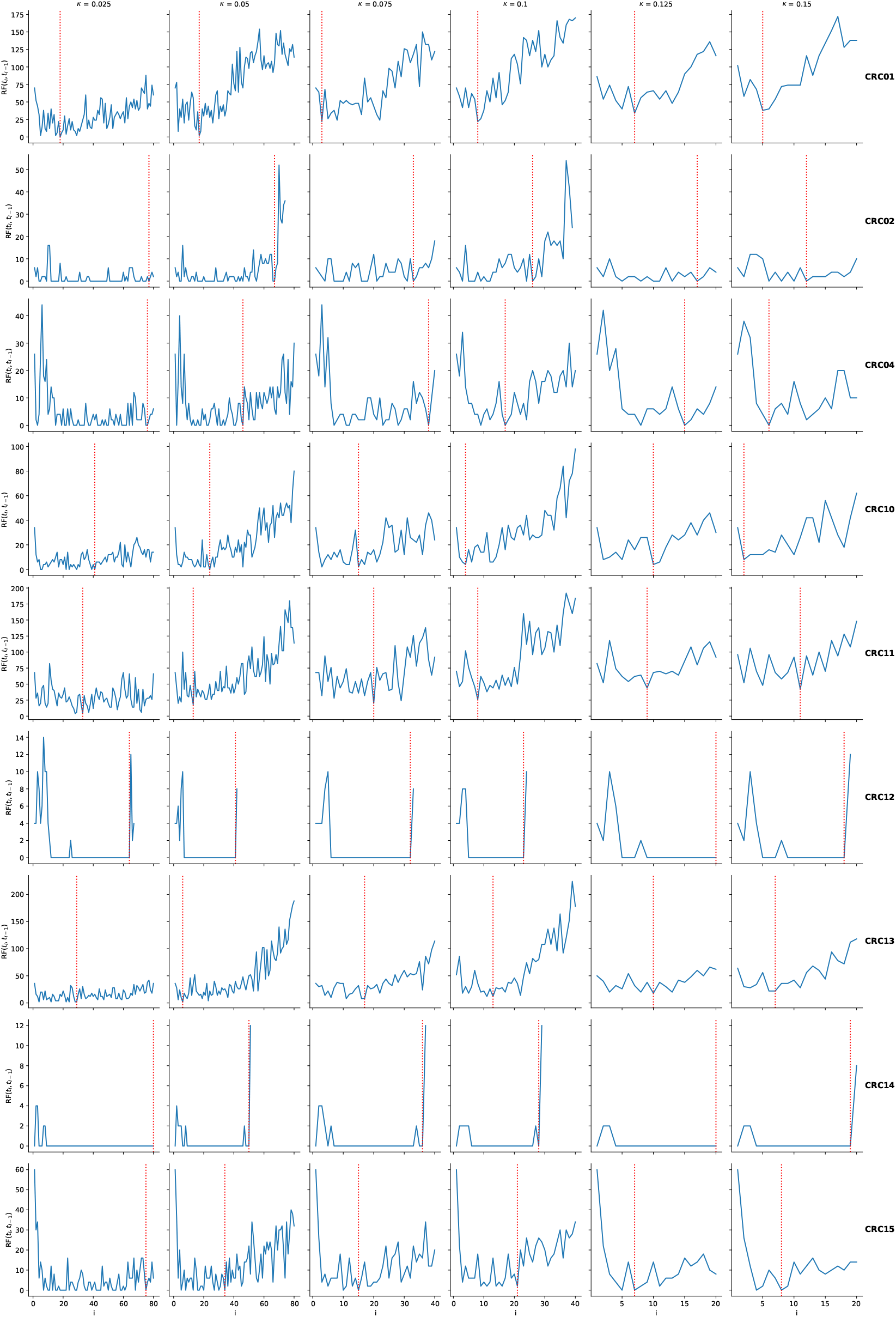
RF distances across iterations for each patient in the Bian *et al*. metastatic CRC cohort applying Sgootr with different *κ* values. RF distances across iterations for each patient in the Bian *et al*. metastatic CRC cohort with different *κ* values. Each row represents each of the 9 patients, and each column represents *κ*=.025, .05, .075, .1, .125, and .15. For *κ*=.025, .05, the iterative procedure was run for a maximum of 80 rounds; for *κ* = .075, .1, maximum 40 rounds; and for *κ*=.125,.15, maximum 20 rounds. Red vertical lines marks the iteration `Sgootr` identifies as optimal. If the blue line ends before the maximum iteration, it means at the last iteration of pruning, the distance between some cells pairs can no longer be computed as all their shared CpG sites have been pruned out. Note that the maximum iteration is arbitrarily chosen, and `Sgootr` does not need to finish running the maximum iteration in order to detect whether the optimal condition has been met.

**Fig. S10:**
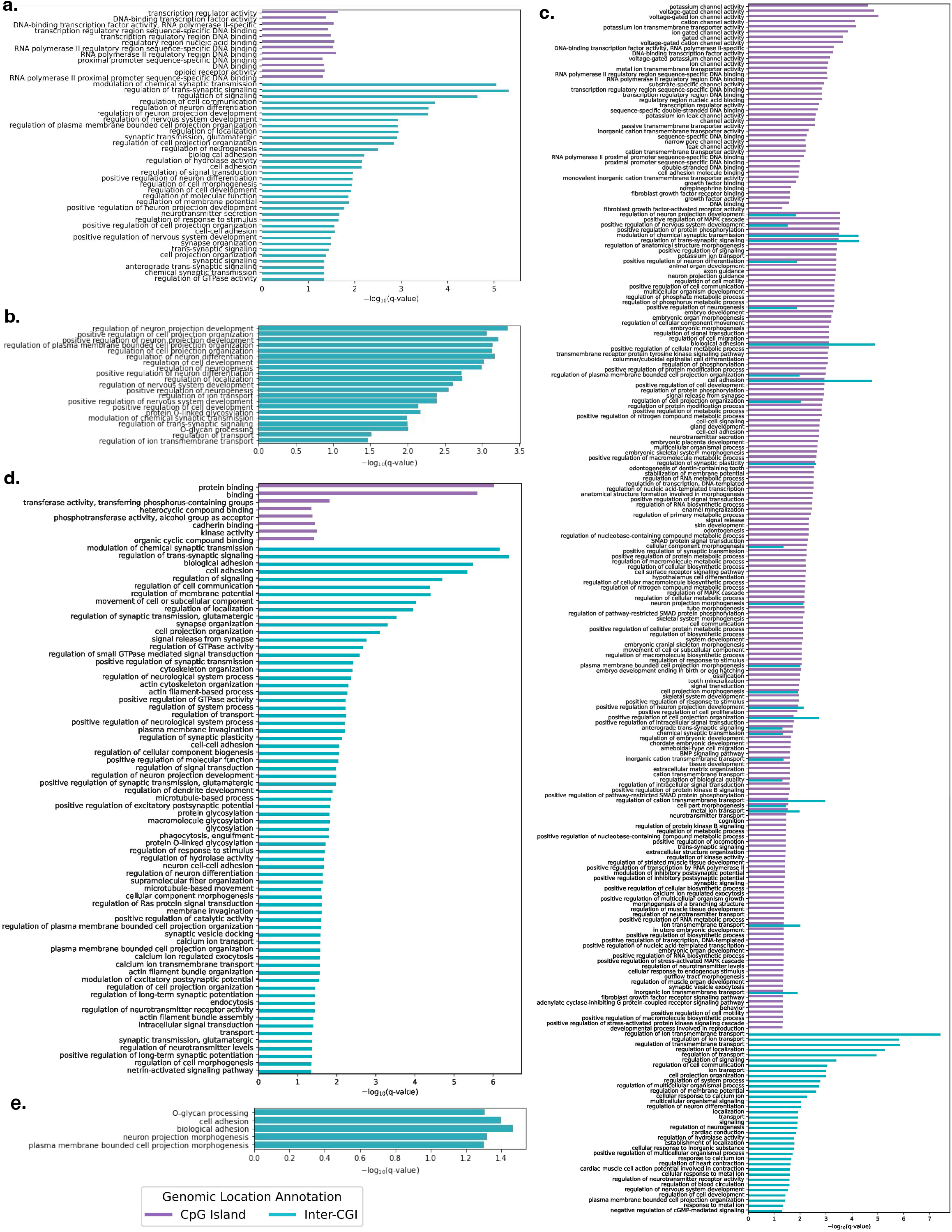
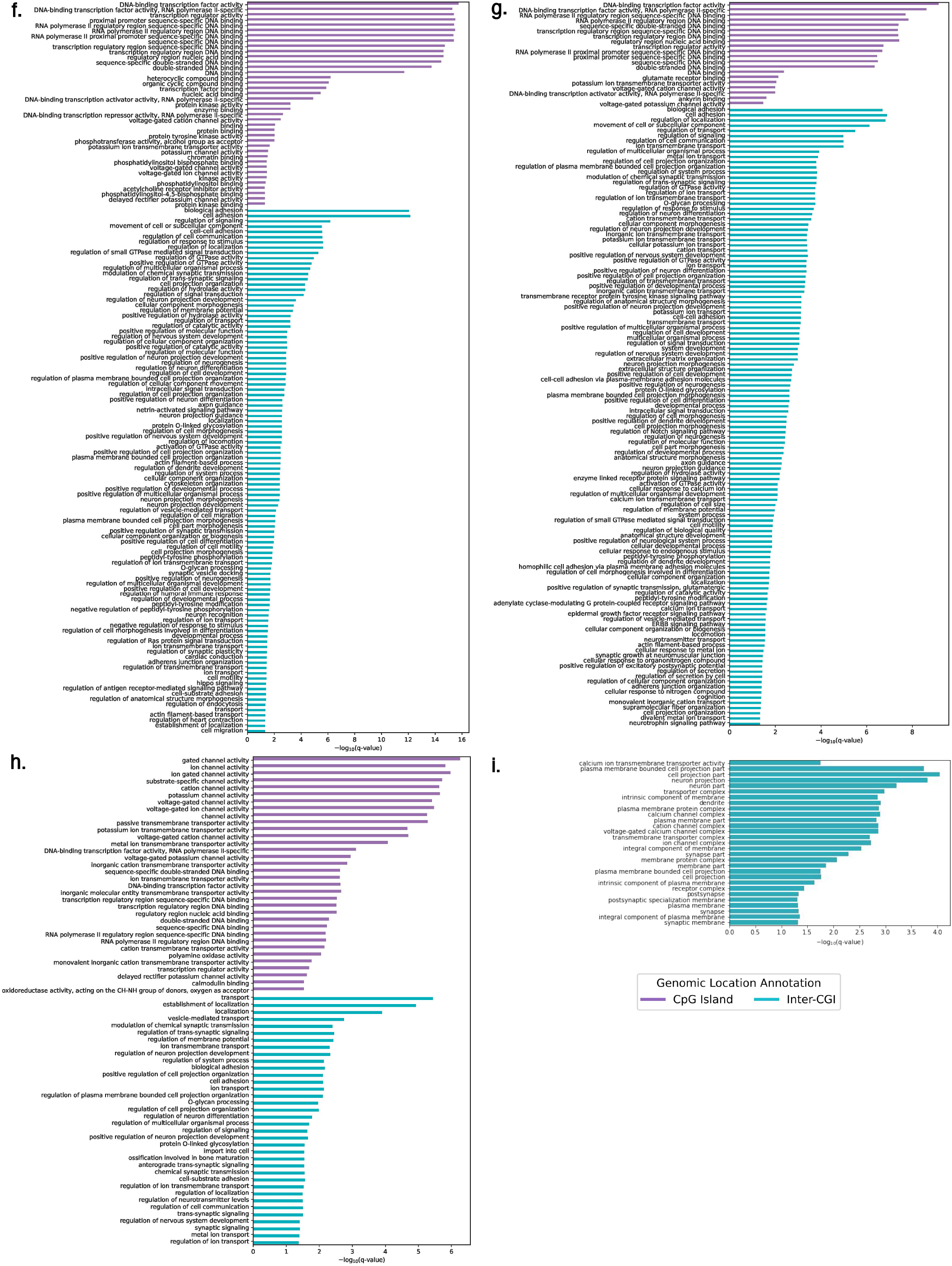
GOrilla[11]-identified GO terms with significant (<.05) q-values in enrichment analysis of non-pseudo, protein coding genes whose gene bodies and 1000bp upstream promoter regions contain the `Sgootr`-identified lineage-informative CpG sites, stratified by whether they locate in CpG island or inter-CGI regions for a. CRC01, b. CRC02, c. CRC04, d. CRC10, e. CRC13, f. CRC11, g. CRC12, h. CRC14, and i. CRC15.

**Fig. S11:**
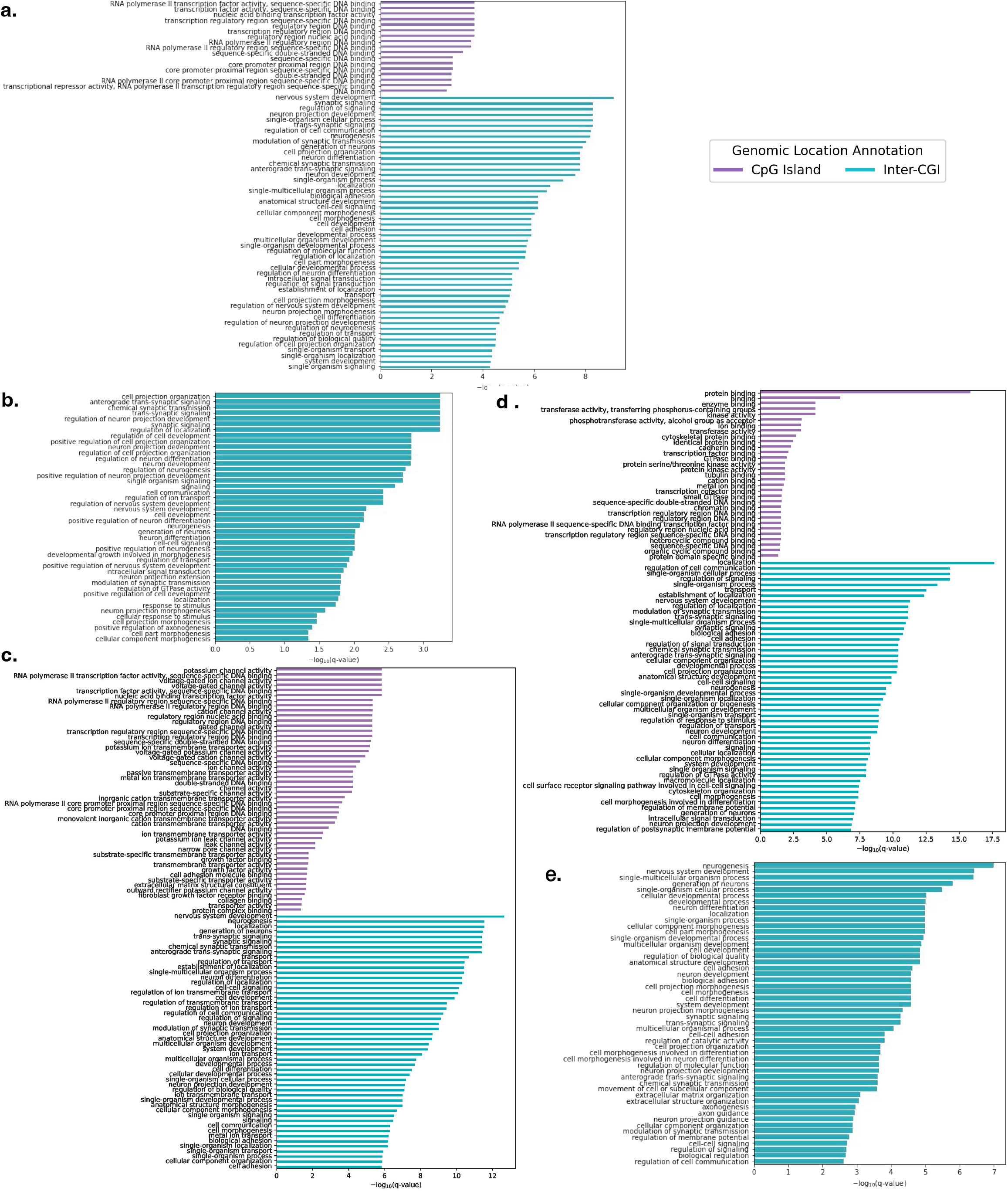
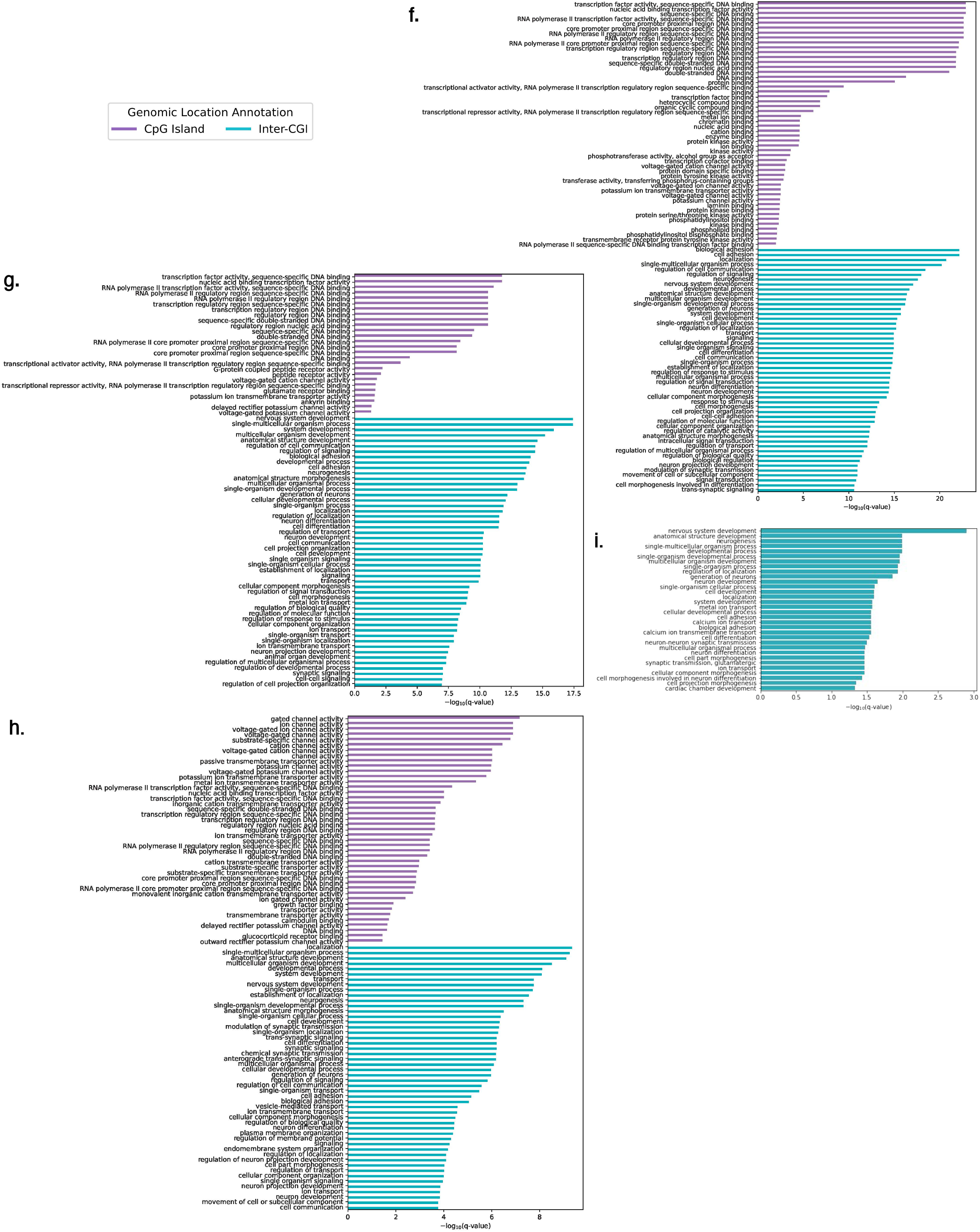
DAVID[36, 18]-identified GO terms with significant (<.05) q-values in enrichment analysis of non-pseudo, protein coding genes whose gene bodies and 1000bp upstream promoter regions contain the `Sgootr`-identified lineage-informative CpG sites, stratified by whether they locate in CpG island or inter-CGI regions for a. CRC01, b. CRC02, c. CRC04, d. CRC10, e. CRC13, f. CRC11, g. CRC12, h. CRC14, and i. CRC15.

**Fig. S12:**
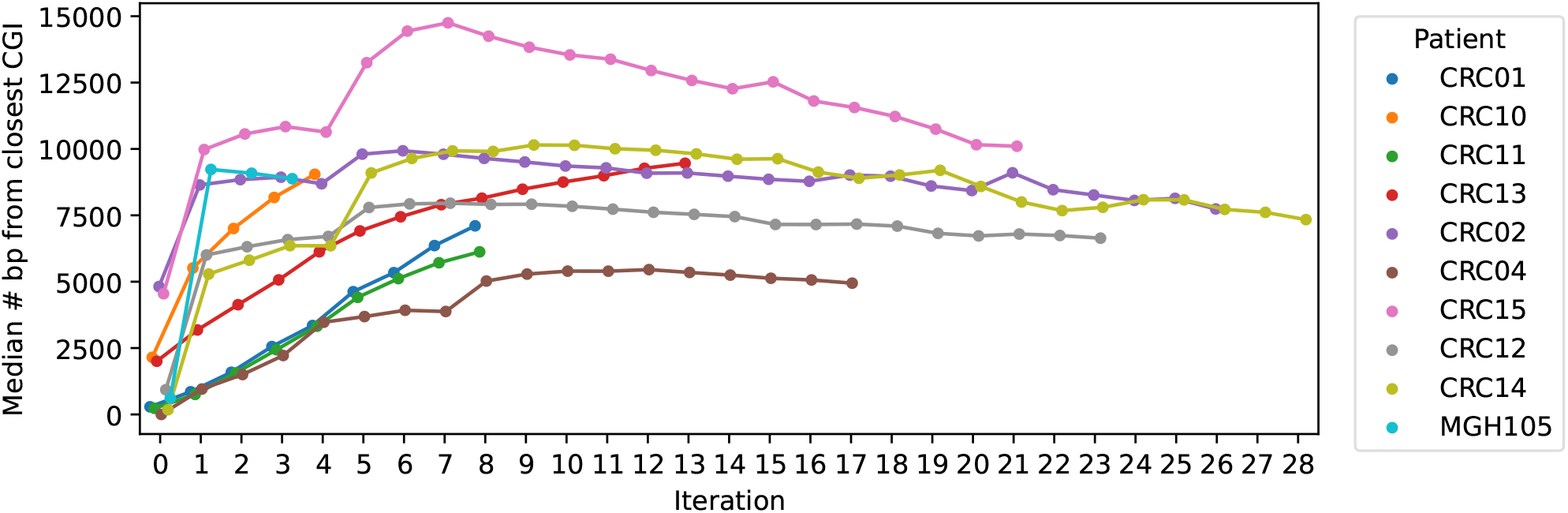
Median number of bp to the closest CGI from CpG sites used to construct the methylation lineage tree at each iteration for all CRC patients and GBM patient MGH105. If a CpG site is within an CGI, it is 0 bp away from the closest CGI.

**Table S4:** Overlap coefficients of GO terms associated with different genomic locations (CpG island and inter-CGI) discovered by GOrilla and DAVID. CRC02, CRC13, and CRC15 don’t have any terms discovered by either GOrilla or DAVID in CpG island. The GO terms associated with inter-CGI regions in CRC15 discovered by GOrilla and DAVID belong to different sub-ontologies (molecular function and biological process, respectively), and are thus not comparable, despite having common themes in neural activities (Figure S10i. and S11i.).

| Patient | CpG Island | Inter-CGI |
| --- | --- | --- |
| CRC01 | 0.8182 | 0.9412 |
| CRC02 | NA | 0.7620 |
| CRC04 | 0.7500 | 0.8689 |
| CRC10 | 1.0000 | 0.8571 |
| CRC11 | 0.9730 | 0.9200 |
| CRC12 | 0.95 | 0.9160 |
| CRC13 | NA | 0.6 |
| CRC14 | 0.8824 | 0.8333 |
| CRC15 | NA | NA |

**Table S5:** Spearman correlation of CpG site distance from closest CpG island and homogeneity.

| Patient | Spearman's Rho | p-value |
| --- | --- | --- |
| CRC01 | -0.5977 | 0.0 |
| CRC02 | -0.3006 | 0.0 |
| CRC04 | -0.6224 | 0.0 |
| CRC10 | -0.5827 | 0.0 |
| CRC11 | -0.6369 | 0.0 |
| CRC12 | -0.4700 | 0.0 |
| CRC13 | -0.5354 | 0.0 |
| CRC14 | -0.5922 | 0.0 |
| CRC15 | -0.4431 | 0.0 |
| MGH105 | -0.3907 | 0.0 |

## S4 Application of Sgootr on the Bian *et al*. cohort [2], replacing NJ with FastME 2.0

**Fig. S13:**
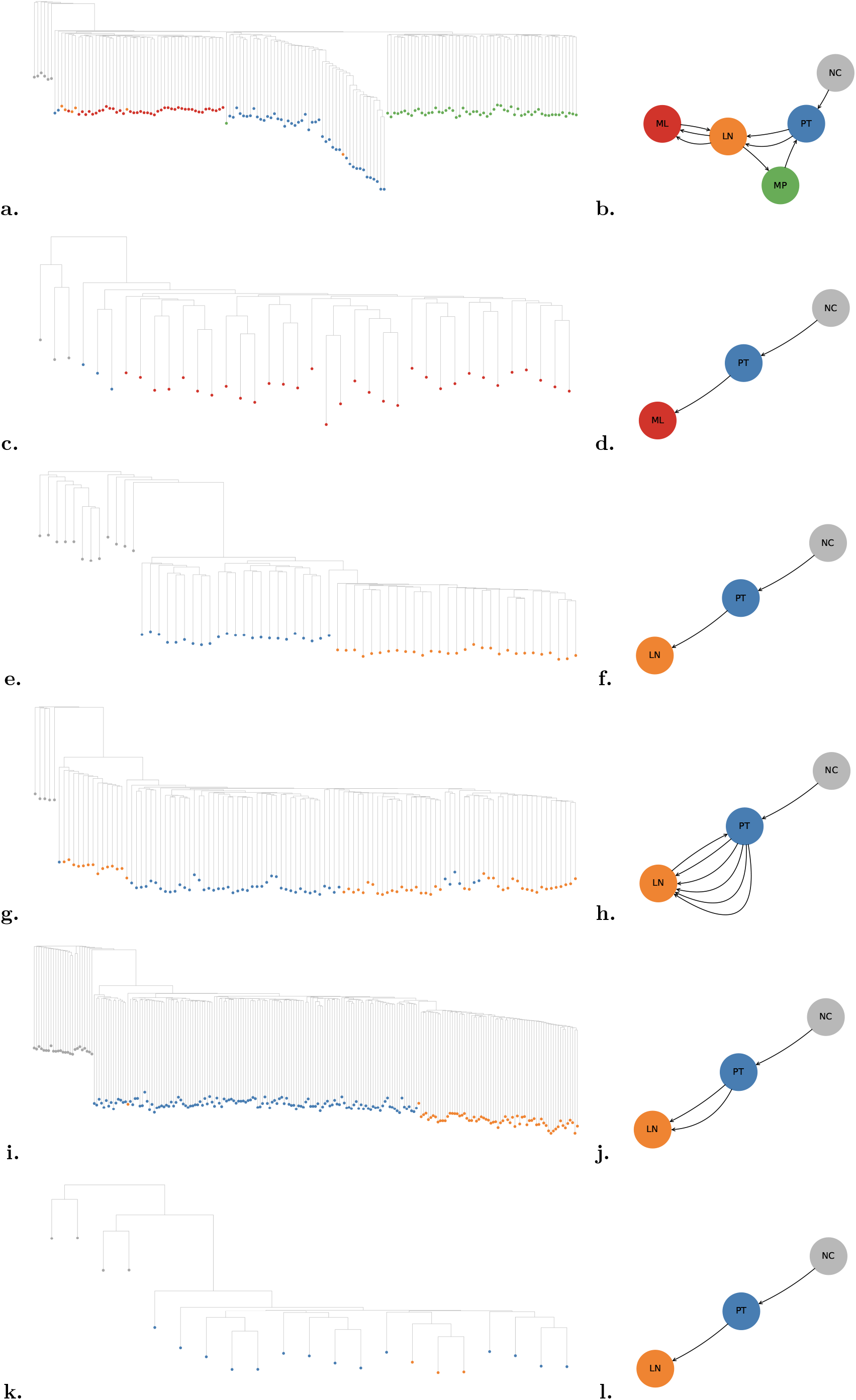
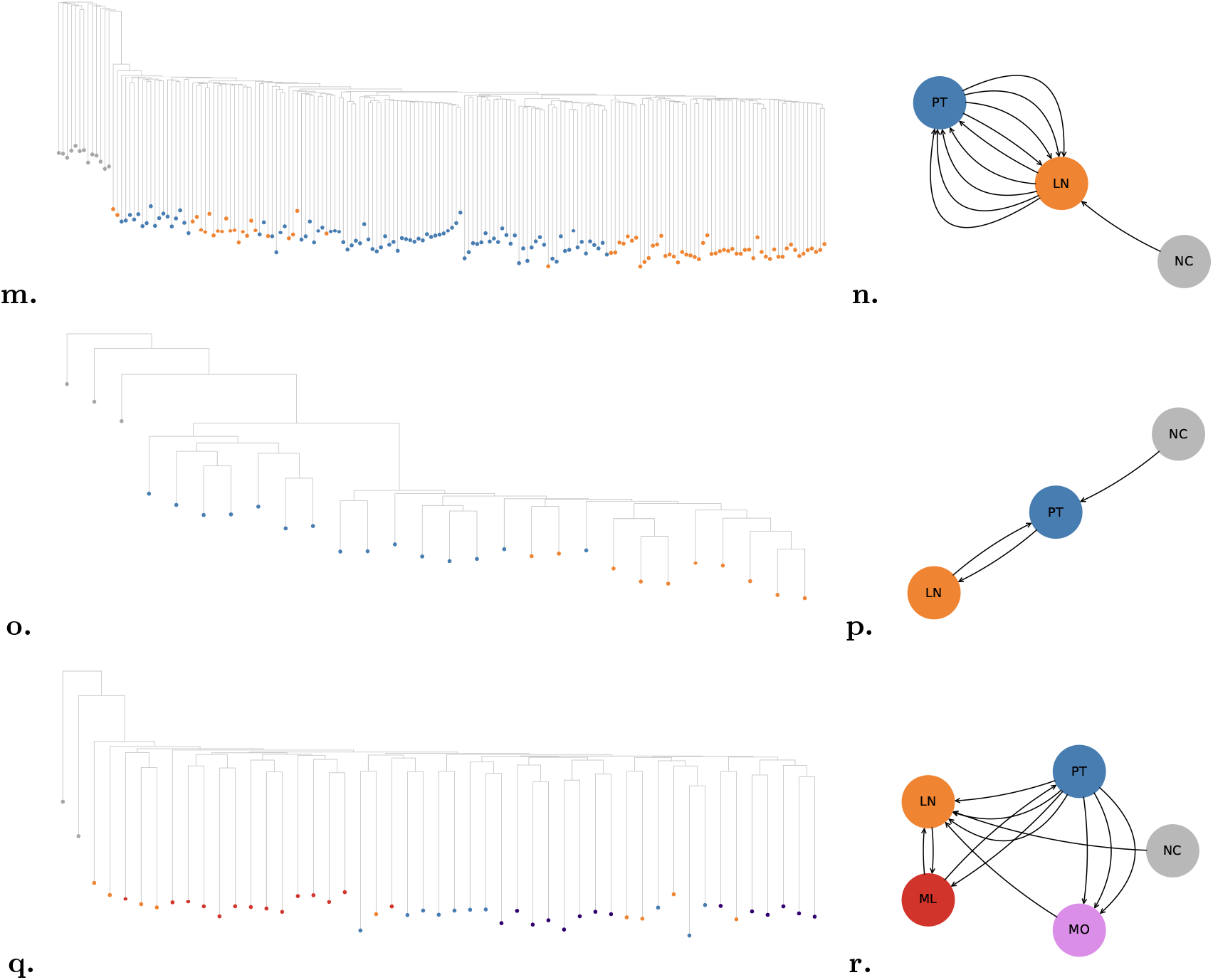
Sgootr-constructed (*κ* = .1) tumor lineage trees (*t*^∗^s) and -inferred tumor migration histories for patients in the Bian *et al*. cohort, replacing NJ with FastME 2.0: CRC01 (a., b.), CRC02 (c., d.), CRC04 (e., f.), CRC10 (g., h.), CRC11 (i., j.), CRC12 (k., l.), CRC13 (m., n.), CRC14 (o., p.), CRC15 (q., r.)

**Fig. S14:**
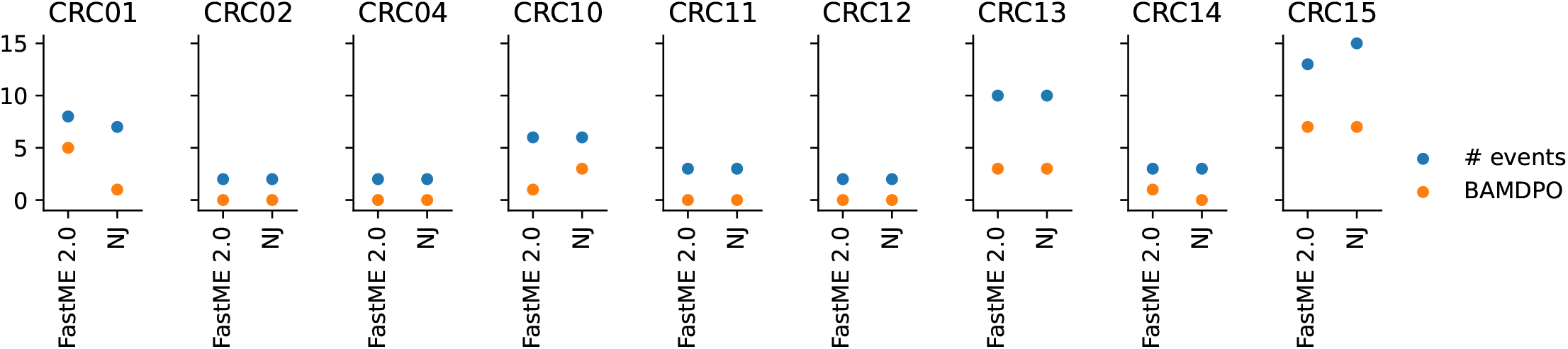
Number of migration events and BAMDPO of migration histories obtained via applying Sgootr with FastME 2.0 in comparison to NJ.

**Fig. S15:**
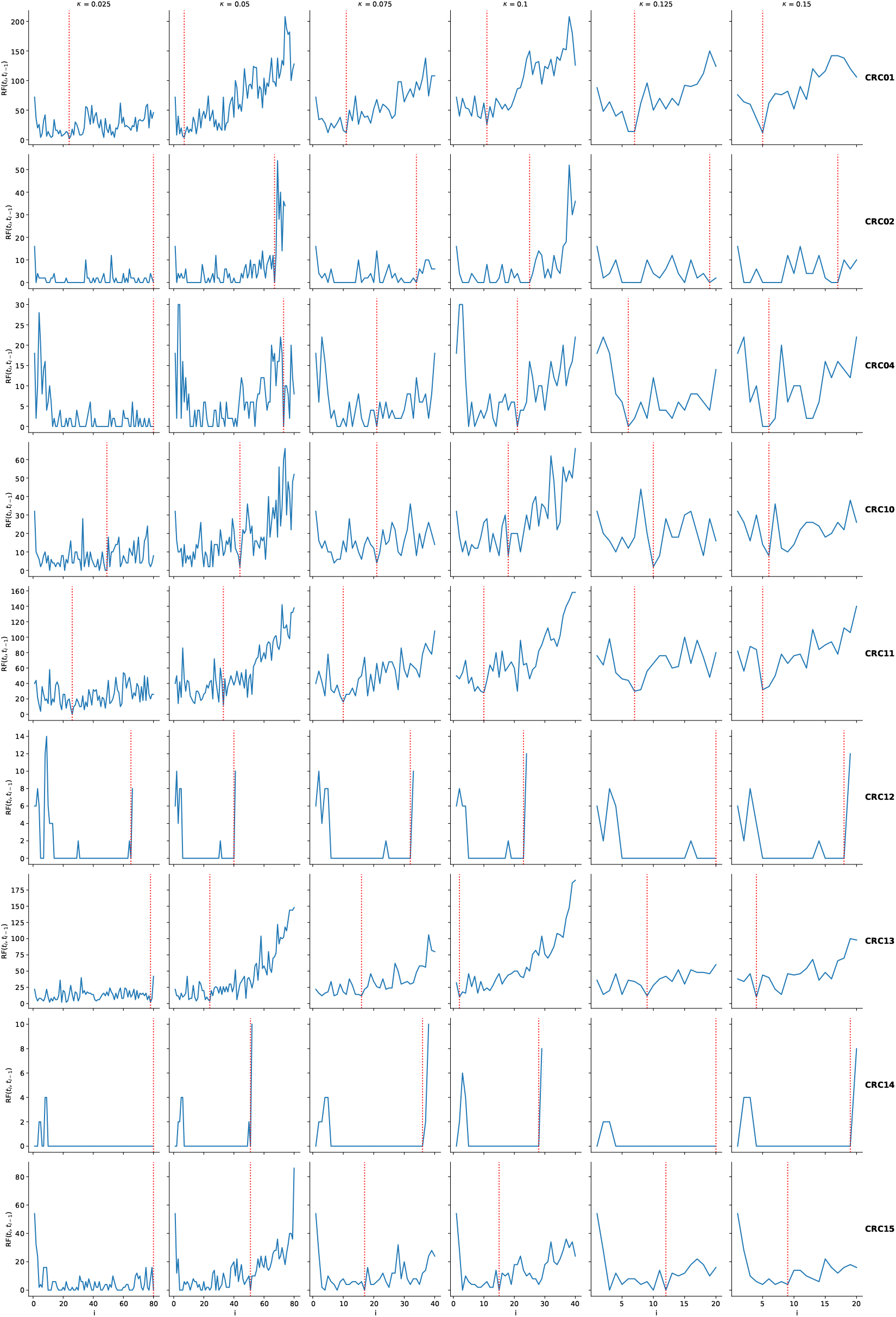
RF distances across iterations for each patient in the Bian *et al*. metastatic CRC cohort applying Sgootr - substituting NJ with FastME 2.0 - with different *κ* values. Each row represents each of the 9 patients, and each column represents *κ*=.025, .05, .075, .1, .125, and .15. For *κ*=.025, .05, the iterative procedure was run for a maximum of 80 rounds; for *κ* = .075, .1, maximum 40 rounds; and for *κ*=.125,.15, maximum 20 rounds. Red vertical lines marks the iteration `Sgootr` identifies as optimal.

**Fig. S16:**
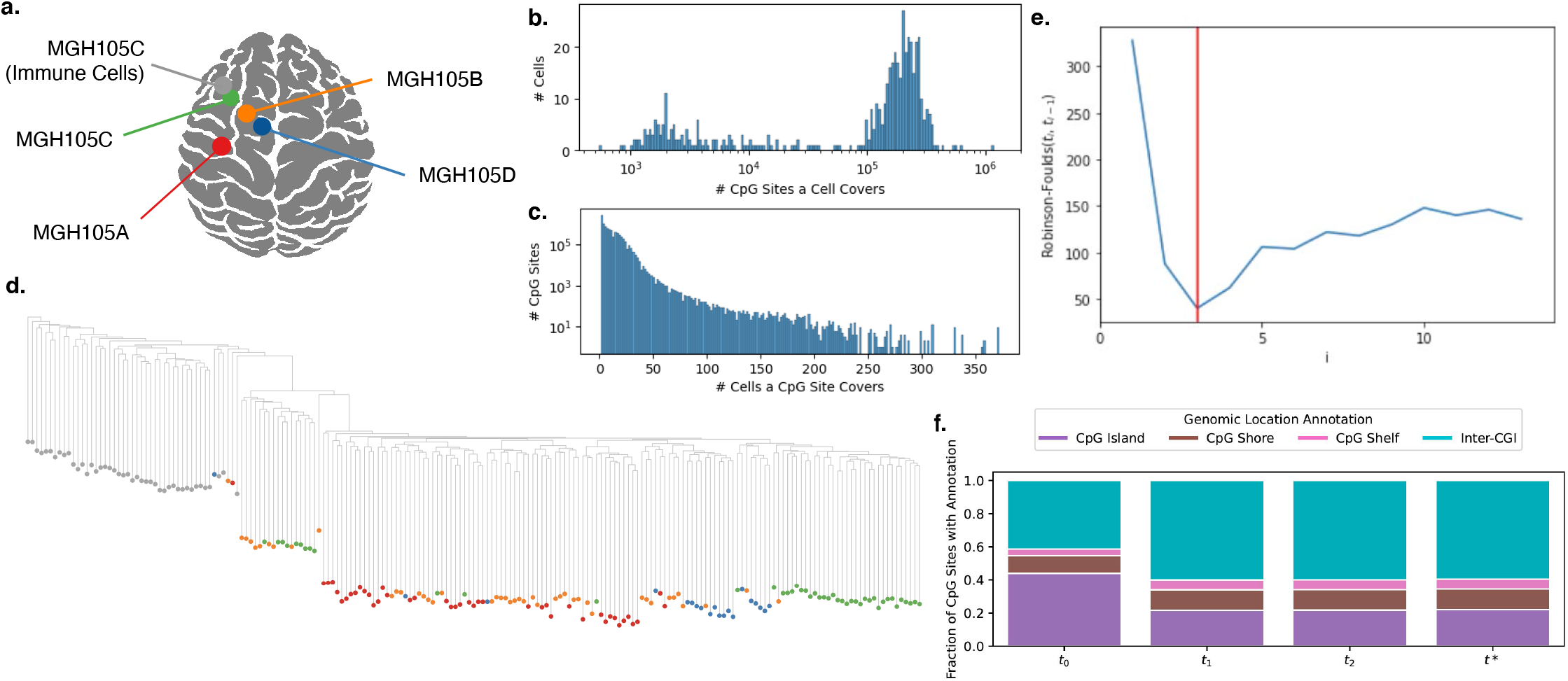
Application of Sgootr to patient MGH105 MscRRBS data by Chaligne *et al*. **a**. The multiregional patient data include single cells from 4 distinct sampling locations (in addition to immune cells from sampling location MGH105C) **b**. Cell coverage distribution among autosomal CpG sites in patient MGH105. Note that the x-axis is in log scale. **c**. Autosomal CpG site coverage distribution in patient MGH105. Note that the y-axis is in log scale. **d**. Tumor lineage tree constructed by `Sgootr` with *κ* = .1 and each single cell at the leaves colored by its sampling location. **e**. RF distances across 14 iterations, with global minimum occurring at *i* = 3, the tree at which is shown above. **f**. Fraction of CpG sites located in CpG island, CpG shore, CpG shelf, and inter-CGI regions in the MGH105 trees at each iteration of the pruning procedure of `Sgootr`, from *t*_0_ to *t*∗.

## S5 Application of Sgootr to multiregional GBM patient MGH105 MscRRBS data by Chaligne *et al*.[5]

We apply `Sgootr` to the MscRRBS data from MGH105, the only multiregionally sampled GBM patient from the Chaligne cohort [5]. Single tumor cells are obtained from 4 sampling locations within the same lesion: MGH105A, MGH105B, MGH105C, and MGH105D; additionally, non-tumor immune cells are sampled from MGH105C (Figure S16). Since both per cell and total CpG coverage in MscRRBS dataset are lower in comparison to scBS-seq (Figure S16b.,c.), we modify the parameters of `Sgootr` such that it: selects cells covering no fewer than 200,000 CpG sites, selects autosomal sites covered in no less than 5% of the remaining cells, bypasses the post-filtering step, and relaxes the pruning parameters so that *δ* = .02, *ω* = .05. We root the trees with an immune cell from sampling location MGH105C.

`Sgootr` obtains the optimal lineage tree *t*∗ = *t*_3_ (Figure S16d.,e.), and the clustering implied by the tree topology seems to highly correspond to the sampling locations of the single cells. We attempted to construct a lineage tree on the same set of cells and sites as we use here, with the set of methods Chaligne *et al*. leveraged in the original work[5], namely FG+IQ-TREE; however, FG fail to terminate within 100 hours hence no result is reported.

## S6 How to choose *p*, P(1^*c*^), P(0^*c*^), and P(mixed) for Sgootr

While in our experiments, we default to *p* = .5, P(1^c^)=P(0^c^)=P(mixed)=.33, it is possible for users to choose other values with additional pieces of prior knowledge. Here we offer examples of when the users may choose other values for those parameters.

The user may set *p >* .5 if there is reasonable belief that, given a pair of alleles truly heterozygous for a CpG site, the sequencing technology will sample from the methylated allele with higher probability than from the unmethylated one. This may manifest in a given dataset as a consistent observation of a higher number of methylated reads than unmethylated reads at diploid heterozygous sites. Similarly, the user may set *p <* .5 if there is reasonable belief that the unmethylated allele is more likely sampled given a heterozygous pair. If strong evidence for such sequencing bias is present, the user should adjust the value of p accordingly.

The user may have P(1^c^)=P(0^c^)»P(mixed), if there is reasonable belief that true heterozygous CpG sites are rare. In this case, at low-coverage CpG sites, the observation of reads of a single status will give diminished probability that the site is truly heterozygous in the expected distance computation. In another potentially extreme example, the user may set P(1^c^)»P(0^c^)=P(mixed), if there is reasonable belief that the homozygously methylated status is prevalent, and homozygously unmethylated and heterozygous statuses are equally rare. In this case, at a particular low-coverage CpG site, heterozygous status will only be given diminished probability if all observed reads are methylated; however, if all observed reads are unmethylated, some larger-than-diminished probability will still be given to the heterozygous status.

## S7 How to choose *κ* for Sgootr

In the main text, we present `Sgootr` results with *κ* = .1. Having also experimented with a range of *κ* values, namely *κ* = {.025, .05, .075, .1, .125, .15} (Figure S9,S15), we provide some guidance on how to choose *κ* for future applications.

Recall that our goal is to detect a point in the iterative procedure where most non-lineage-informative sites have been pruned out but most lineage-informative sites still remain. Translating the objective into measurable terms, we return the last iteration where the global minimum RF distance occurs, after the initial decrease and before the ultimate increase.

When the RF distances across iterations appear flat, or have a region where RF distances have small fluctuations, and iterations where global minimum occur are far from unique - as one can observe across many patients for *κ* = .025, .05, .075 columns in Figure S9,S15 - *κ* is likely too small. With small *κ*, `Sgootr` prunes out fewer sites at each iteration. Intuitively, RF distances between trees constructed from more similar sets of sites will be lower, possibly explaining the over all low RF distances in many instances. In those cases, the turning point in the RF distances becomes difficult, if not impossible, to observe, and a higher *κ* is called for.

While higher *κ* often lead to clear identification of a unique global minimum in RF distances across iterations - as one can observe across many patients for *κ* = .125, .15 columns in Figure S9,S15 - it bares the risk of over-pruning. One may consider attempting lower values of *κ* while the V-shaped RF distance trend can still be observed.

Of course, determining the maximum number of iterations to run goes hand-in-hand with determining *κ*. If resource availability allows, we recommend choosing a number of iterations such that a consistent increase in RF distance towards the later iterations can be observed.

## S8 Experiment details

In this section, we provide details and parameters used to generate the results presented in this work.

### S8.1 Sgootr

While we specifically mention in main text the parameters used to generate the `Sgootr` results presented in this work, see `config.yaml` at the GitHub repository https://github.com/algo-cancer/Sgootr for the full set of default `Sgootr` configurations in one place.

### S8.2 FastME 2.0

In constructing a tumor lineage tree from a distance matrix using FastME 2.0 instead of NJ, we ran FastME 2.0 with the following command: `fastme -i {.mat distance matrix file} -o {.nwk output} -n -s`

### S8.3 IQ-TREE

We ran iqtree with the following command, referencing what was done in prior work by Gaiti *et al*. [14] and Chaligne *et al*. [5]: `iqtree2 -s {.phy file} -st BIN -m TESTNEW -mset GTR2 -nt AUTO -bb 1000 -wbtl -abayes -alrt 1000 -nstop 500 -wsr -nm 2000 -v -alninfo -seed {1, 2, 3, 4, 5}`

### S8.4 FG

For FG, we follow the method descriptions by Gaiti *et al*.[14]. Given the cell-by-site matrices for methylated and unmethylated reads, we first obtain the binarized methylation status for each covered site in a cell via the procedure described in Section 3. We further prune out sites with coverage in less than 5 cells. For each remaining site *s*, we randomly downsample the other sites by 100x, and count the number of times *s* forms four gametes with sites in the downsampled set, disregarding the pair if they are within a read length of each other (150bp for Bian *et al*., and 50bp for Chaligne *et al*.) or do not have at least 4 cells with coverage for both sites. For each site, we divide the number of occurrences of four gametes by the total number of pairings, obtaining the FG frequency.

The sites are binned into intervals of width 0.05 with regard to its methylation rate (number of cells with methylated status over total number of cells with read coverage for that site), disregarding bins with rates <0.1 or >0.9, and for each bin compute the median FG ratio. Within each bin, sites whose FG frequency is 1.5x median absolute deviation away from the bin median frequency were selected. The aggregate set of sites across all bins form the output of FG.

While random downsampling is necessary for problems of the scale in this work, it also introduces stochasticity. Like Gaiti *et al*., we perform 5 separate runs of FG for each experiment, and found high concordance of the per site FG frequencies across the runs (Figure S18). For each site, we compute the mean FG frequency across 5 runs to use for the selection process described above (Figure S17).

**Fig. S17:**
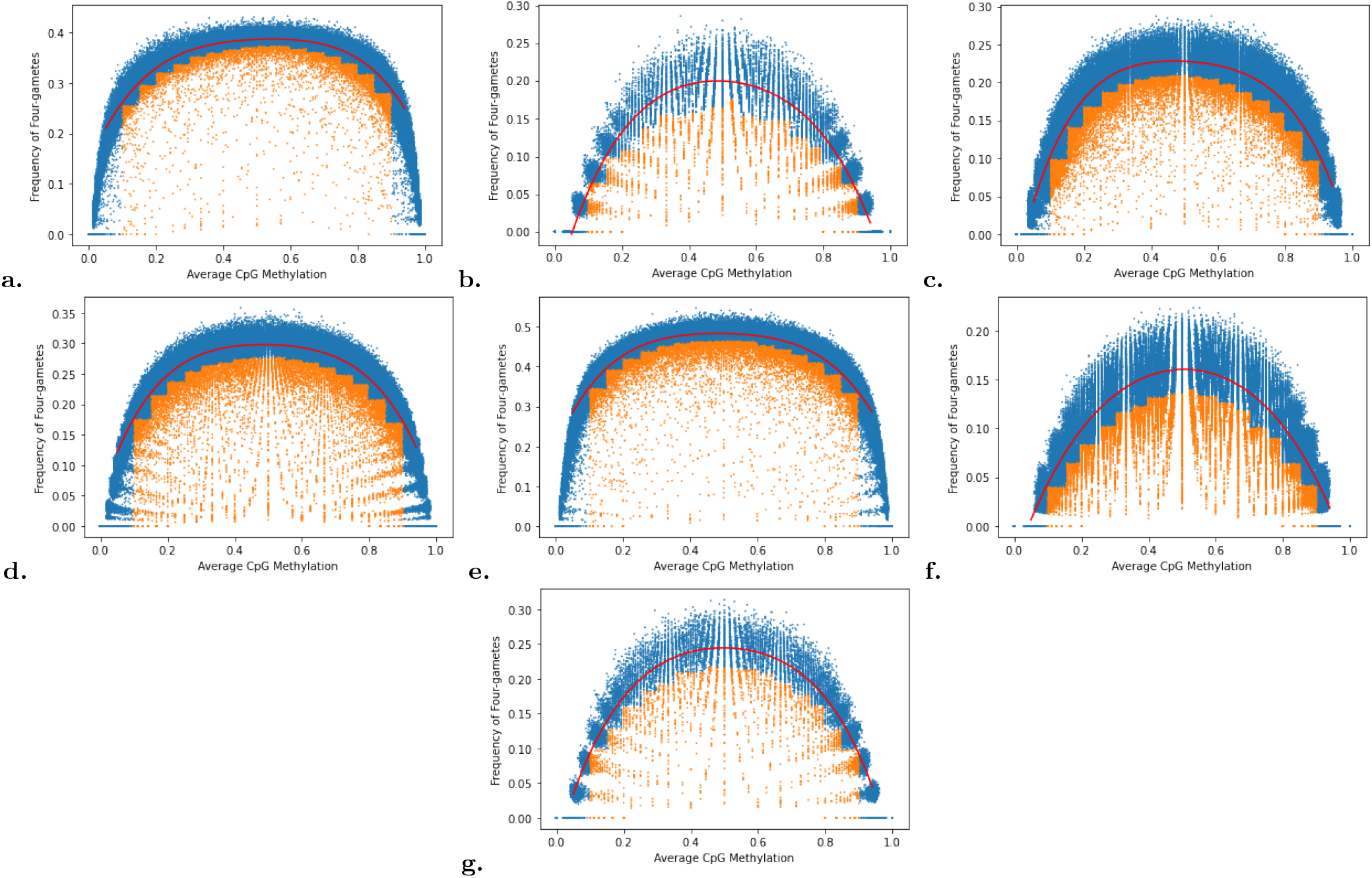
FG frequencies (averaged across 5 runs) for a. CRC01, b. CRC02, c. CRC04, d. CRC10, e. CRC13, f. CRC14, g. CRC15. Selected sites are colored with orange. The red line denote the smoothed line connecting the median FG frequency within each bin.

**Fig. S18:**
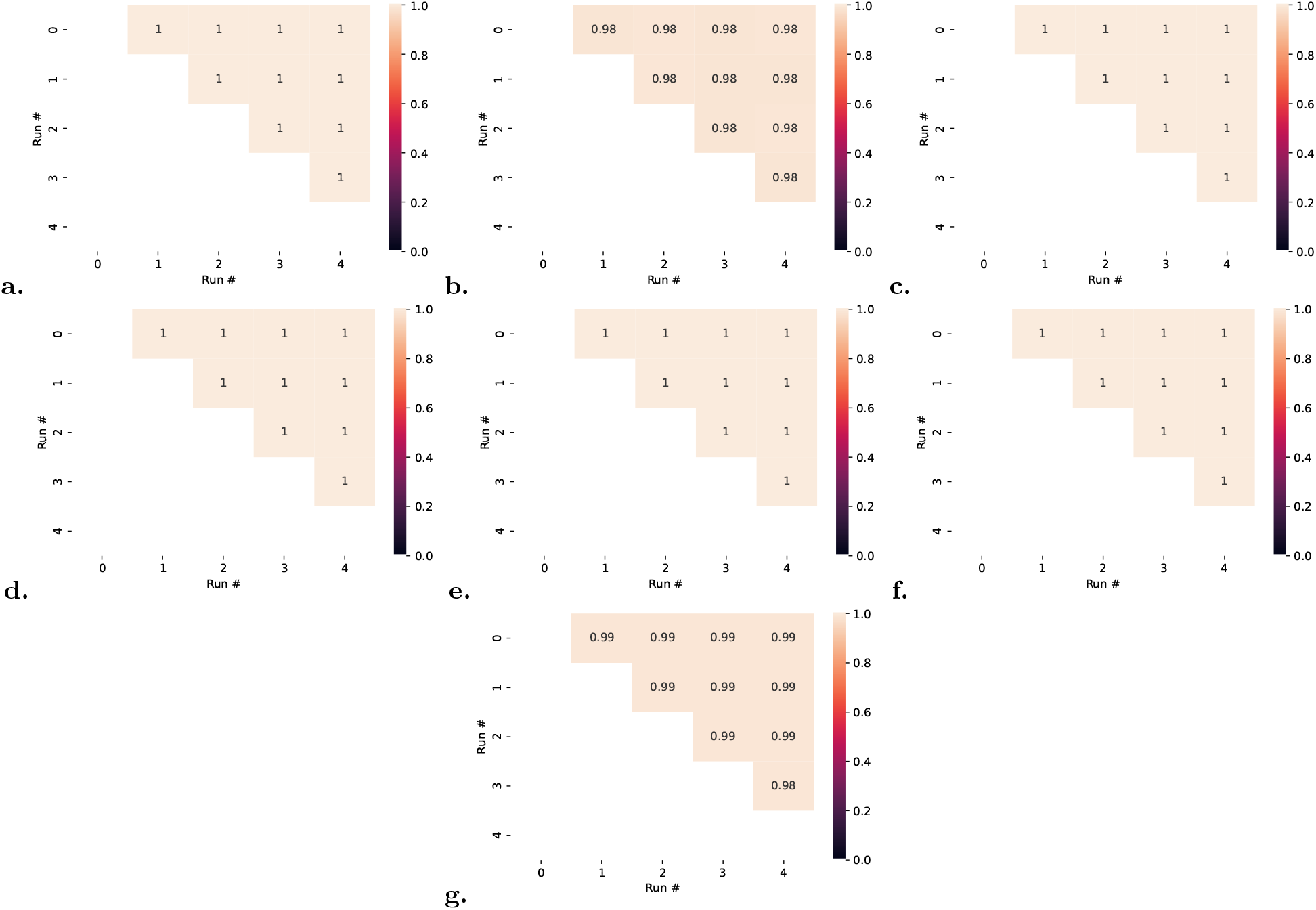
Pearson product-moment correlation coefficients between FG frequencies across sites between down-sampled runs for a. CRC01, b. CRC02, c. CRC04, d. CRC10, e. CRC13, f. CRC14, g. CRC15. Results from each pair of runs from a single experiment exhibit high concordance.

## Footnotes

8 Out of 12 total patients in the Bian *et al*. cohort, we excluded those without scBS-seq data (CRC03 and CRC06), and those without metastasis information (CRC09).

9 from the Trinity CTAT Project https://github.com/broadinstitute/inferCNV

